# A Pillar and Perfusion Plate Platform for Robust Human Organoid Culture and Analysis

**DOI:** 10.1101/2023.03.11.532210

**Authors:** Soo-Yeon Kang, Masaki Kimura, Sunil Shrestha, Phillip Lewis, Sangjoon Lee, Yuqi Cai, Pranav Joshi, Prabha Acharya, Jiafeng Liu, Yong Yang, J. Guillermo Sanchez, Sriramya Ayyagari, Eben Alsberg, James M. Wells, Takanori Takebe, Moo-Yeal Lee

## Abstract

Human organoids have potential to revolutionize *in vitro* disease modeling by providing multicellular architecture and function that are similar to those *in vivo*. This innovative and evolving technology, however, still suffers from assay throughput and reproducibility to enable high-throughput screening (HTS) of compounds due to cumbersome organoid differentiation processes and difficulty in scale-up and quality control. Using organoids for HTS is further challenged by lack of easy-to-use fluidic systems that are compatible with relatively large organoids. Here, we overcome these challenges by engineering “microarray three-dimensional (3D) bioprinting” technology and associated pillar and perfusion plates for human organoid culture and analysis. High-precision, high-throughput stem cell printing and encapsulation techniques were demonstrated on a pillar plate, which was coupled with a complementary deep well plate and a perfusion well plate for static and dynamic organoid culture. Bioprinted cells and spheroids in hydrogels were differentiated into liver and intestine organoids for *in situ* functional assays. The pillar/perfusion plates are compatible with standard 384-well plates and HTS equipment, and thus may be easily adopted in current drug discovery efforts.

## Introductiond

Recent advances in successful establishment of various culture protocols have led to differentiation of pluripotent stem cells (PSCs) into brain, lung, liver, stomach, intestine, kidney, and pancreas organoids^1,2^. While human organoids represent a new direction in predictive *in vitro* toxicity screening, there are several technical challenges to adopt human organoids in preclinical evaluations of drug candidates^3^. Current organoid cultures rely on the ability of PSCs to self-organize into discrete tissue structures with a step-wise process that mimics normal organ development^4^. This allows for a closer approximation to the structural and functional complexity of adult tissues, and therefore more accurate modeling of development and disease. Although organoid transplantation into animals can induce vascularization and enhance growth and maturation, insufficient maturity of organoids cultured *in vitro* remains the biggest challenge in organoid research. Strategies to recreate structural complexity focus on the directed differentiation of PSCs into organ tissues by manipulating signaling pathways in a temporal manner that correspond to organogenesis^5,6^. To better model diseases, next generation organoid systems have been developed, including liver organoids with inflammatory cells^7^ or intestinal organoids with enteric nerves^8^ or fluidic coupling of multiple tissue modules^9^.

Despite the potential of organoids to model human physiology and disease, they have been surprisingly underutilized for high-throughput screening (HTS) of compounds. This is in part because there is a lack of miniature organoid culture systems that are compatible with HTS equipment. Conventionally, organoid differentiation and maturation processes have been performed in 6-/24-well plates, petri dishes, and spinner flasks, which are impractical for HTS of compounds and highly labor-intensive mainly due to a large volume of expensive cell culture media and growth factor cocktails required and repeated dissociation and encapsulation of organoids in hydrogels necessary to control organoid size and density. There is still a lack of miniature, easy to use, high-throughput culture systems that allow small-scale organoid culture for HTS. Recently, suspension of human induced PSCs (iPSCs) have been dispensed in an extracellular matrix (ECM)-coated 384-well plate using an automated liquid handling machine and differentiated and matured into kidney organoids for HTS^10^. This approach may not work for many organoid systems due to the necessity of encapsulation of cells in biomimetic hydrogels at different stages of organoid formation and cell death in the core of large organoids. In addition, microfluidic devices have been used recently to culture kidney and brain organoids to enhance maturity^11,12^, but their use for organoid culture is still limited due to low throughput and lack of user friendliness. Labor-intensive, manual PSC and organoid loading into a microfluidic device is a big challenge to facilitate HTS of compounds. In addition, the microchannels in current microfluidic devices are too narrow to culture organoids that are typically 200 μm – 2 mm in diameter, and thus microchambers have to be used for organoid culture. Other 3D cell culture platforms, including ultralow attachment (ULA) well plates, hanging droplet plates, and microcavity array plates, may be unsuited for long-term organoid culture due to the necrotic core from static cell culture and difficulty in accommodating hydrogels in culture. Organoids that are too large (typically bigger than 500 μm) become hypoxic and necrotic due to lack of oxygen and nutrients in the core of organoids^13,14^. Thus, it is essential to control the size of organoids by physical and enzymatic dissociation for reproducible generation of data from organoids. In addition, existing methods of analyzing organoids are still low throughput and do not allow for dynamic monitoring of organoid function, which is a major obstacle to conduct rapid drug screening *in vitro*. Currently, structural and cellular characterization of organoids typically require fixing or harvesting the organoid for immunostaining, which are low-throughput and do not measure dynamic changes in real time^15^. Constant monitoring of cultures as they progress through an experimental treatment allows for observation of immediate responses (e.g., action potential firing, dynamic hormone secretion) as well as long-term changes (e.g., time lapse imaging of morphological changes). Live imaging of organoids may enable continuous monitoring of flow-perfused organoids^16^ as well as those expressing fluorescent reporters^8,17^.

These issues are a significant bottleneck for successful implementation of organoids for high-throughput, predictive compound screening. To address some of these issues, we have developed unique microarray three-dimensional (3D) bioprinting technology and associated pillar/perfusion plates, including a 384-pillar plate with sidewalls and slits (“384PillarPlate”) and a clear-bottom, 384-deep well plate (“384DeepWellPlate”) for static organoid culture as well as a 36-pillar plate with sidewalls and slits (“36PillarPlate”) and a 36-perfusion well plate with reservoirs and microchannels (“36PerfusionPlate”) for dynamic organoid culture. Microarray 3D bioprinting is a microsolenoid valve-driven, high-precision, robotic cell printing technology demonstrated on pillar/perfusion plates to create human organoids rapidly and reproducibly with minimal manual intervention (**Fig. 1 and Suppl. Fig. 1**). In the present study, the pillar/perfusion plates have been successfully manufactured *via* injection molding of polystyrene, which is non-cytotoxic and minimizes nonspecific adsorption of compounds. Highly reproducible human liver organoids (HLOs) and human intestine organoids (HIOs) have been created by precisely dispensing foregut cells suspended in the mixture of alginate and Matrigel and transferring mid-hindgut cell spheroids in Matrigel on the pillar plates and then differentiating and maturing them in highly controlled oxygen/nutrient diffusion environments, which led to replication of miniature tissues *in vitro* with critical liver and intestine functions. In summary, we envision that bioprinted human organoids in the pillar/perfusion plate system could be used for high-throughput, predictive human toxicology and disease modeling.

**Figure 1.**
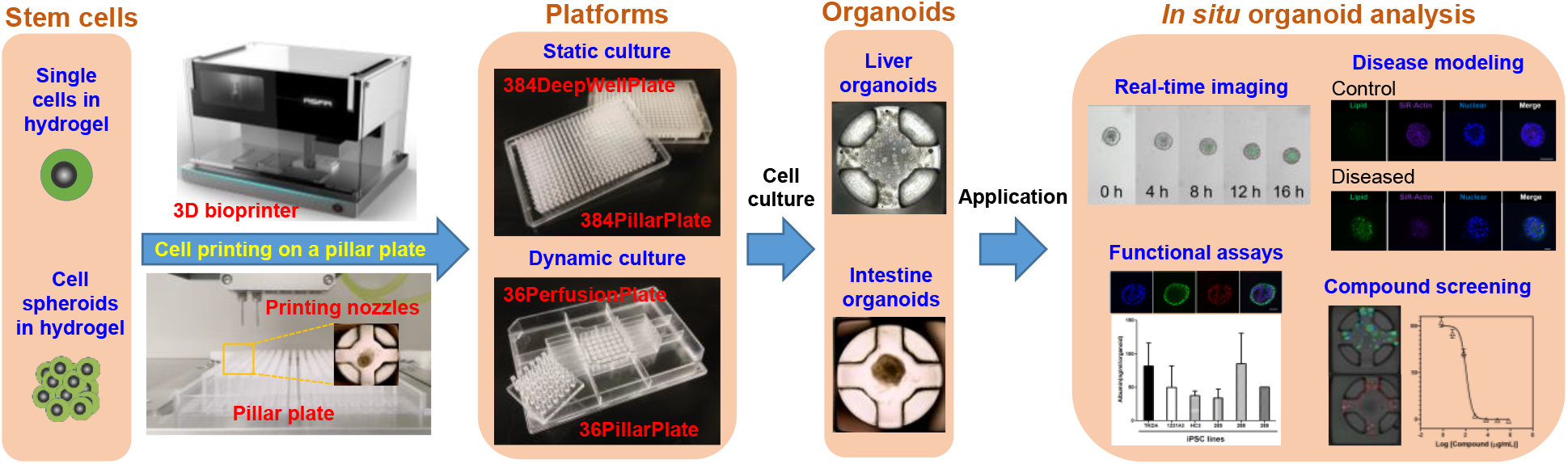
Microarray 3D bioprinting technology and associated pillar and perfusion plate platform for human organoid culture and analysis. Pluripotent stem cells (PSCs) and spheroids suspended in biomimetic hydrogels can be printed on the pillar plates precisely with a 3D bioprinter. After hydrogel gelation, the pillar plates containing PSCs can be sandwiched with a deep well plate or a perfusion well plate with differentiation and maturation media for static and/or dynamic cultures of organoids. Bioprinted organoids can be tested with compounds, stained with fluorescent dyes and antibodies, and scanned with an automated fluorescence microscope for high-content imaging (HCI) of organoid functions as well as predictive assessment of compound toxicity.

## Results

### Development of microarray 3D bioprinting technology and associated pillar and perfusion plates

Scaling up organoid production for HTS would require increasing the numbers of tissue culture plates or transitioning cultures to spinner flasks, and then dispensing organoids into a HTS system^18^. Alternatively, PSCs can be dispensed directly into a HTS system and differentiated into human organoids *in situ*. The former case is easier for HTS assays when mature organoids are “small” (below 100 μm in diameter) and uniform in size and shape for consistent dispensing in hydrogels using automated liquid handling machines. Except for tumor organoids derived from biopsy samples, mature organoids are typically 200 μm – 2 mm in diameter, difficult to dispense robustly into high-density well plates, and hard to passage using proteolytic and collagenolytic enzymes such as Accutase^®^ due to low cell viability after dissociation. Therefore, mechanical dissociation methods are commonly introduced for passaging mature organoids, which may lead to organoids of irregular size and difficulty in robotic dispensing. The latter case is more robust in terms of dispensing PSCs, but less convenient for HTS assays due to difficulty in changing growth media frequently without disturbing cells in hydrogels in high-density well plates. Since there are limited cell culture systems developed to address this unmet need, we developed microarray 3D bioprinting technology and associated pillar/perfusion plates (**Fig. 1 and Suppl. Fig. 1**). Our pillar/perfusion plate platform combining “3D bioprinting” with “microfluidic-like” features offers several distinctive advantages over conventional static 3D cell culture models and dynamic microphysiological systems (MPSs). The pillar plates enable dispensing of cells and spheroids suspended in biomimetic hydrogels *via* microarray 3D bioprinting and provide a high-throughput capability by controlling individual pillars with organoids. Multiple cell arrays demonstrated in microfluidic chambers in MPSs typically cannot be controlled individually, resulting in low throughput^19^. The system presented here is highly flexible so that organoids on the pillar plates can be cultured statically and tested dynamically (or *vice versa*) by simply separating the pillar plate and sandwiching onto the deep well plate or the perfusion well plate. In addition, the perfusion well plate allows connection of multiple organoid types on the pillar plate without using micropumps or tubes, which is a critical feature for modeling complex diseases. Furthermore, the pillar/perfusion plates are compatible with standard 384-well plates and existing HTS equipment such as automated fluorescence microscopes and microtiter well plate readers already available in laboratories. Bioprinted organoids with key organ functions can be analyzed on the pillar/perfusion plate by *in situ* tissue clearing and high-throughput, high-content cell imaging. Thus, there is high potential for adopting the pillar/perfusion plate platform with human organoids for predictive compound screening.

A highly versatile 384PillarPlate with sidewalls and slits and a complementary 384DeepWellPlate have been manufactured *via* injection molding of polystyrene to support organoid cultures. A single 384PillarPlate contains 384 pillars onto which an array of human iPSCs suspended in hydrogels has been dispensed using a 3D bioprinter or a multichannel pipette (**Fig. 2 and Suppl. Fig. 1**). Prior to injection molding, various shapes and structures of pillars with sidewalls and slits have been tested *via* 3D printing of transparent plastics for optimum cell printing, formation of a gel for cell encapsulation, and analysis of cells in different layers for high-content imaging (HCI) assays for miniature tissue regeneration. The 384PillarPlate with 4.5 mm of the pillar-to-pillar distance and 11.6 mm of the pillar height has been manufactured by injection molding (**Fig. 2A**). The unique sidewall and slit structure of the 384PillarPlate ensured more robust cell spot attachment for HCI and immunofluorescence assays as compared to the pillars with a flat surface^20^. In addition, proprietary, hydrophilic surface functionalization of the pillar plates has been performed to avoid air bubble entrapment on top of the pillars and retain cells in hydrogels over long time periods (typically 4 weeks). Each pillar can accommodate an optimum volume of 5 μL of hydrogel containing cells without bubble entrapment and a maximum volume of 7 μL without spillage. Polystyrene used for injection molding of the 384PillarPlate is nontoxic for cell culture and transparent for image acquisition of organoids. The 384DeepWellPlate built on a footprint of conventional 384-well plates has 384 deep wells and is complementary to the 384PillarPlate to accommodate growth media for cell culture and reagents for cell staining. The 384DeepWellPlate with 4.5 mm of the well-to-well distance and 3.5, 3.5, and 14.7 mm of the width, length, and depth of wells has been manufactured by injection molding (**Fig. 2B**). Each deep well can accommodate a maximum solution volume of 120 μL, and an optimum volume of 80 μL after sandwiching with the 384PillarPlate for static organoid culture without the overflow. By sandwiching the 384PillarPlate onto the 384DeepWellPlate, biochemical and cell-based assays can be performed in the plate system.

**Figure 2.**
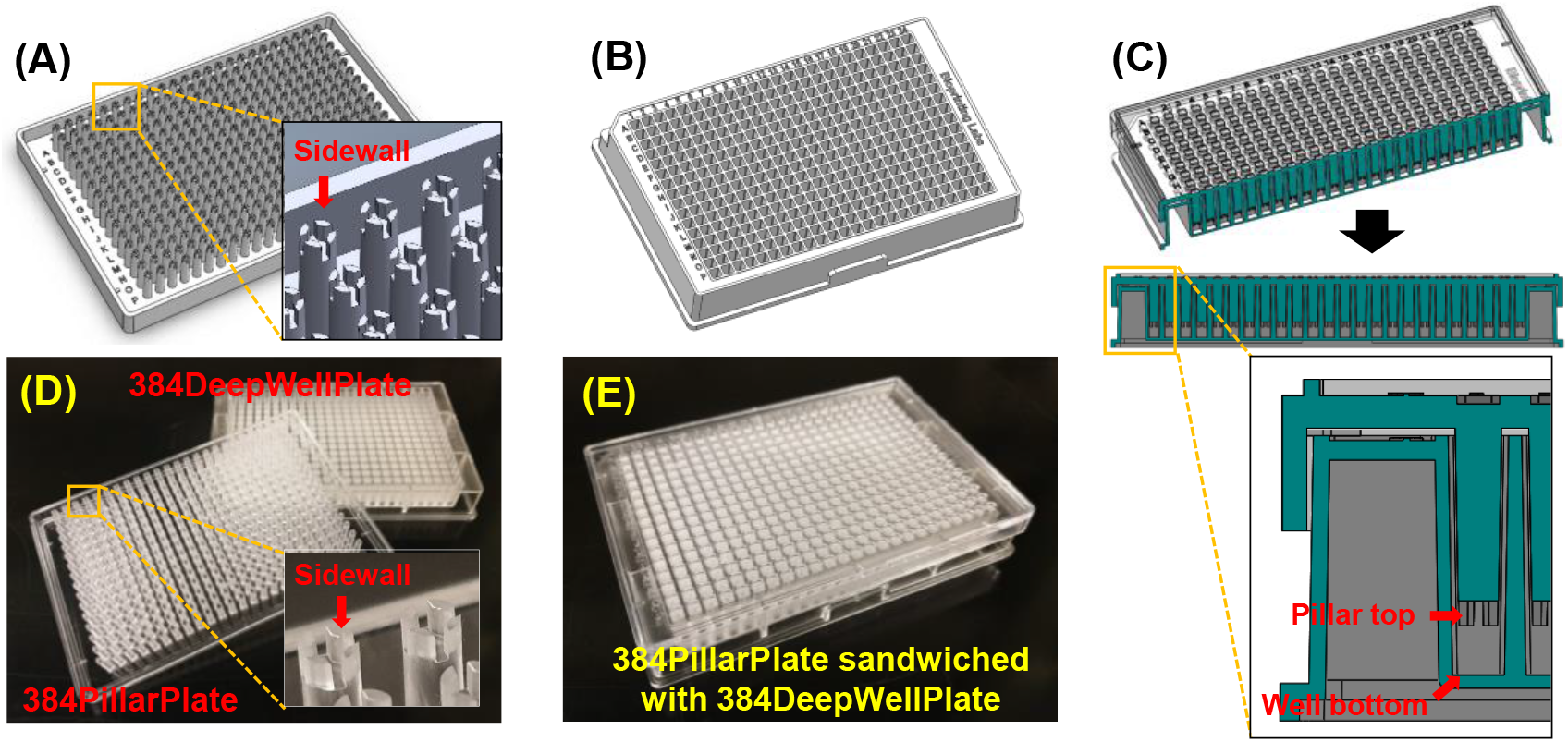
The combination of a pillar plate and a deep well plate for static organoid culture. SolidWorks designs of **(A)** a 384PillarPlate, **(B)** a 384DeepWellPlate, and **(C)** the 384PillarPlate sandwiched onto the 384DeepWellPlate. Sidewalls and slits on the pillars are designed for stem cell printing and robust spot attachment for long-term static organoid culture. Injection molding with polystyrene was performed to manufacture **(D, E)** the 384PillarPlate and the 384DeepWellPlate for static organoid culture.

In addition to the 384PillarPlate and the 384DeepWellPlate, a 36PillarPlate and a 36PerfusionPlate were built on the footprint of conventional 384-well plates by injection molding with polystyrene for dynamic organoid culture (**Fig. 3**). A single 36PillarPlate contains 36 pillars onto which an array of human organoids can be cultured in biomimetic hydrogels. The dimensions of the 36PillarPlate are identical to that of the 384PillarPlate except for the size of the plate and the number of pillars. The 36PerfusionPlate complementary to the 36PillarPlate contains up to 1200 μL of growth media in each of twelve reservoirs and typically 50 - 80 μL of solution in 36 perfusion wells. A thin polystyrene film was attached at the bottom of the 36PerfusionPlate to create microchannels, which connect reservoirs and perfusion wells for dynamic organoid culture and organoid communication. Each channel consists of one upper reservoir (UR), one lower reservoir (LR), and six perfusion wells. To control flow rates within microchannels in the 36PerfusionPlate, it was necessary to change the surface property from hydrophobic to hydrophilic using robust surface coating with proprietary polymer solutions, leading to uniform flow rates within microchannels (**Figs. 3F-G**). There was no flow generated in the 36PerfusionPlate without surface coating due to high surface tension of the hydrophobic polystyrene. Hydrophilic surface coating was important to minimize air bubble entrapment in microchannels, which is critical for uniform organoid culture within the 36PillarPlate. By sandwiching the 36PillarPlate onto the 36PerfusionPlate, biochemical and cell-based assays can be performed with unidirectional or bidirectional flows generated on a digital rocker (**Suppl. Fig. 3**). The optimal volume of growth media within each channel was approximately 800 – 1,000 μL to avoid an “overflow” after sandwiching and rocking. For dynamic organoid culture, the perfusion plate can be tilted on a digital rocker for removing old growth media from reservoirs and replacing with fresh growth media (**Suppl. Fig. 3E**). Velocity profiles in microchannels and perfusion wells were simulated using SolidWorks software (**Figs. 3F-G**). The transport phenomenon considered in this model was the Navier-Stokes equation for free laminar flow. The geometry of the rectangular perfusion wells was designed with two reservoirs. The model assumed that the 36PillarPlate was sandwiched onto the 36PerfusionPlate. By changing the volume of water in the reservoirs, tilting angles on a digital rocker, and frequency of angle change, we were able to determine that the flow rate within the microchannels and the velocity profiles within the perfusion wells to avoid diffusion limitation of growth media on the 36 pillars. The changes in water levels in perfusion wells and reservoirs in the 36PerfusionPlate sandwiched with the 36PillarPlate over time were simulated to avoid overflow (**Fig. 3F**). The bidirectional fluid flow in each channel generated the height difference in reservoirs while maintaining uniform heights within perfusion wells over time. The average flow rate obtained for 900 μL of water at 10° tilting angle and 1 min frequency from the simulation was 6.30 μL/sec, which was comparable to 6.44 ± 0.25 μL/sec from the experiment. Interestingly, high velocity profiles were found under the pillars in the perfusion wells, potentially inducing high fluid mixing under the pillars and accelerating cell growth on the pillars (**Fig. 3G**).

**Figure 3.**
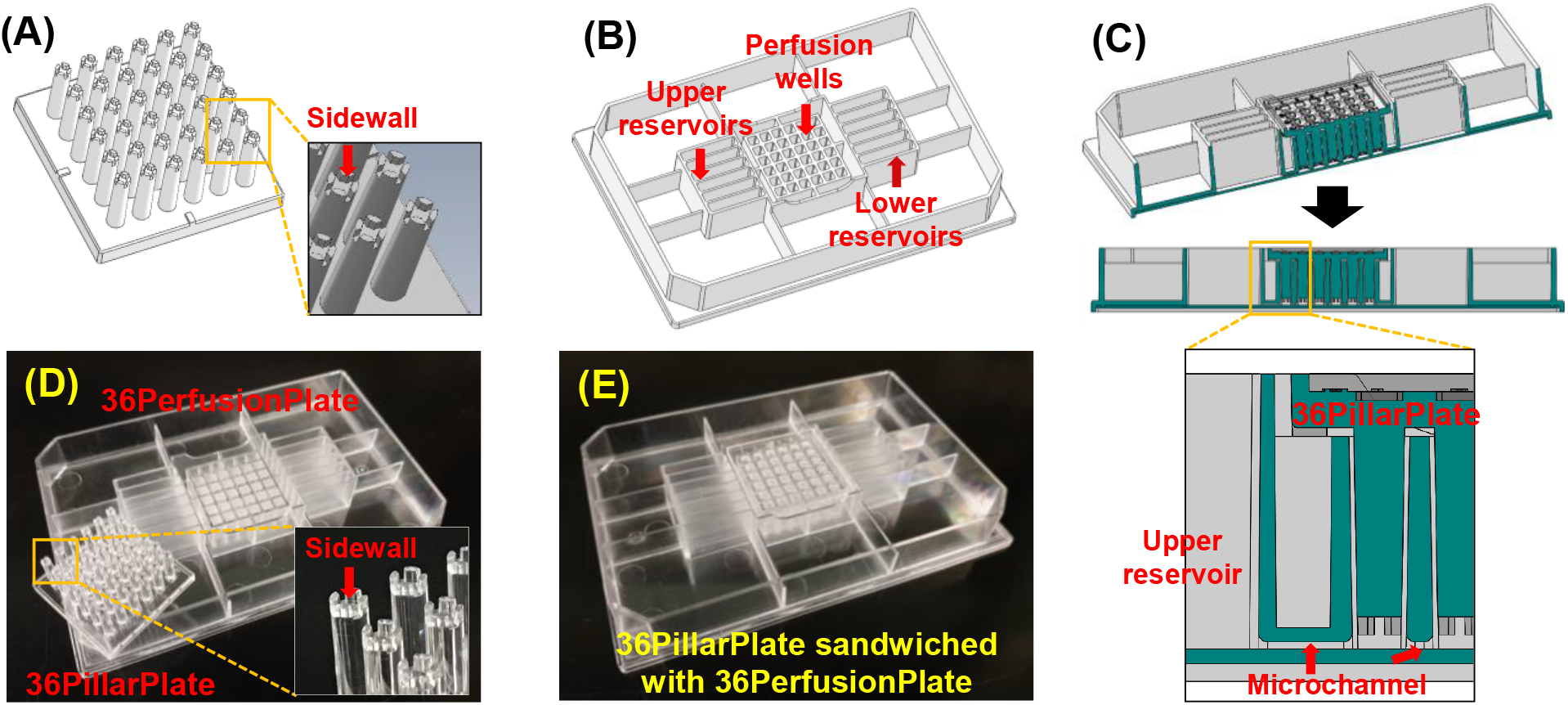
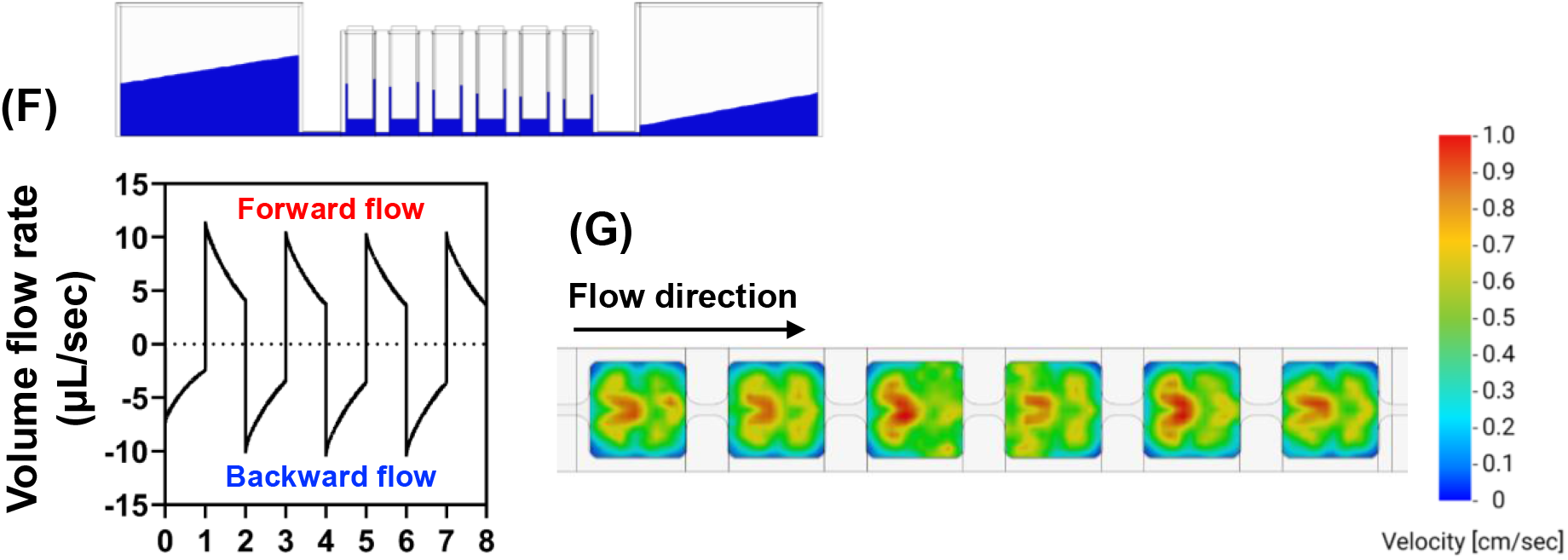
The combination of a pillar plate and a perfusion well plate for dynamic organoid culture. SolidWorks designs of **(A)** a 36PillarPlate, **(B)** a 36PerfusionPlate, and **(C)** the 36PillarPlate sandwiched onto the 36PerfusionPlate. Injection molding with polystyrene was performed to manufacture **(D, E)** the 36PillarPlate and the 36PerfusionPlate for dynamic organoid culture. The culture media added into the upper reservoirs (URs) can flow through the microchannels and the perfusion wells to reach the lower reservoirs (LRs) by gravity. **(F)** Forward and backward flow rates in microchannels of the 36PerfusionPlate on a digital rocker obtained from SolidWorks simulation with 900 μL of water at 10° tilting angle and 1 min frequency. **(G)** Velocity profile under the pillars in the 36PerfusionPlate simulated with SolidWorks with 900 μL of water at 10° tilting angle and 1 min frequency. The surface roughness of the 36PerfusionPlate was equal to a Petri dish (0.0017 μm).

### Rapid and robust cell printing on the pillar plate for human liver organoid (HLO) culture

To establish differentiation of iPSCs into organoids and demonstrate their function on the pillar/perfusion plate platform successfully, the choice of iPSCs at different differentiation stages, biomimetic hydrogels, and growth media with specific cocktails of growth factors and additives are crucial. As a proof of concept, we demonstrated differentiation of human liver organoids (HLOs) and human intestine organoids (HIOs) from foregut cell suspension in the mixture of alginate and Matrigel printed on the pillar plate and mid-hindgut cell spheroids in Matrigel on the pillar plate, respectively. Briefly, frozen foregut cells suspended in the mixture of alginate and Matrigel (**Fig. 4**) or 6-8 mg/mL Matrigel (**Suppl. Figs. 1C-D**) were successfully printed on the pillar plate (5 μL/pillar; 3,000 cells/pillar) using a 3D bioprinter (ASFA™ spotter from MBD Korea) with a chilled printing head equipped with two microsolenoid valves and two custom-made 600 μm orifice nozzles. Only 3 minutes were needed to print foregut cells in Matrigel on all 384 pillars. The printed foregut cells in hydrogels were gelled by sandwiching the pillar plate onto the 384DeepWellPlate and incubating them in a 5% CO_2_ incubator at 37°C for 20 minutes. This is followed by incubating the foregut cells in organoid formation medium with CEPT cocktail (50 nM <u>C</u>hroman 1, 5 μM <u>E</u>mricasan, 1,000-fold diluted <u>P</u>olyamine supplement, and 0.7 μM <u>T</u>rans-ISRIB) in the 384DeepWellPlate and then culturing them in hepatic canaliculi formation medium and hepatocyte maturation medium to mimic the liver differentiation process (**Suppl. Fig. 2A**)^21^. The CEPT cocktail was used instead of Y-27632 for 4 days to recover foregut cells from cryopreservation damage. Morphologically, HLOs cultured both on the pillar plate and in a 24-well plate look similar with a circular epithelial cell layer with internal lumen surrounded by mesenchymal cells (**Figs. 4A-B**). The size of the matured HLOs generated was 0.1 – 0.2 mm. Interestingly, there was mesenchymal cell growth after retinoic acid (RA) treatment in Matrigel (control), which was reduced significantly in the mixture of alginate and Matrigel (**Fig. 4C**). The reproducibility of cell printing on the pillar plate was determined by measuring cell viability after 1 day of cell printing with CellTiter-Glo^®^ luminescent cell viability assay kit and calculating the coefficient of variation (CV). From three trials, high reproducibility of single cell printing in hydrogels was achieved with CV values between 11.8% - 18.6% (**Suppl. Fig. 1C**).

**Figure 4.**
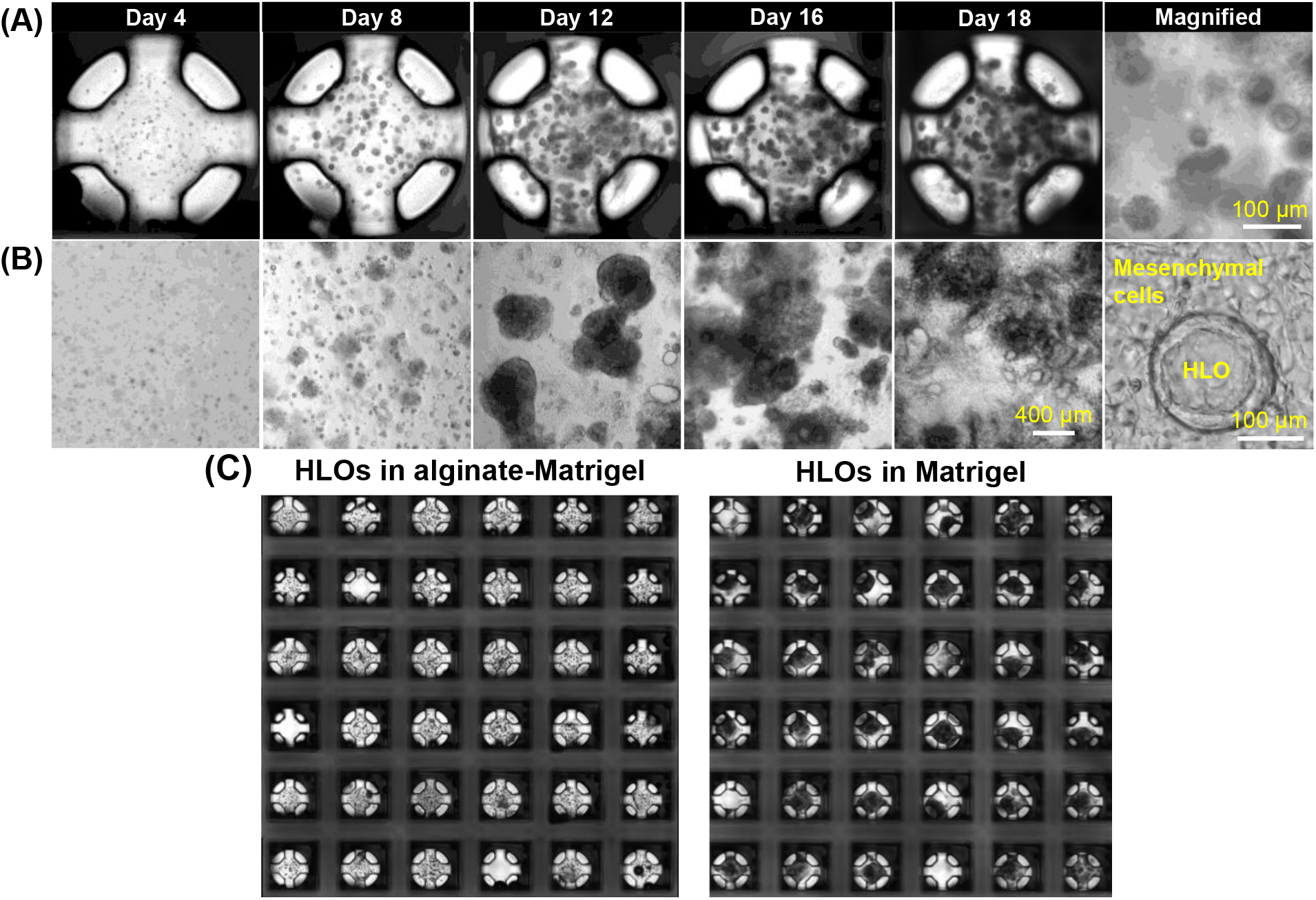
Bioprinted human liver organoids (HLOs) on the pillar plate. **(A)** Bioprinted frozen foregut cells (72-3 cells) encapsulated in the mixture of 0.75% (w/v) alginate and 6-8 mg/mL Matrigel on the 36PillarPlate cultured in the presence of CEPT cocktail for four days, differentiated, and matured into HLOs for three weeks. **(B)** Frozen foregut cells in 2-fold diluted Matrigel domes cultured in a 24-well plate as a control. Viability of frozen foregut cells and spheroid formation were improved significantly with CEPT cocktail. The size of the matured HLOs was 0.1 – 0.2 mm. **(C)** Uniform generation of human liver organoids (HLOs) from 72-3 cells on the 36PillarPlate by seeding 3,000 cells/pillar in the mixture of alginate and Matrigel (Left) or in Matrigel (Right) and differentiated for 18 days on the pillar plate. Over 90% of cell spots were robustly attached on the pillar plate during the long-term culture. There was mesenchymal cell growth after retinoic acid (RA) treatment in Matrigel (control), which was reduced significantly in the mixture of alginate and Matrigel.

### Functionality of static cultured HLOs on the pillar plate

To validate high reproducibility of HLO formation, albumin secretion from HLOs differentiated from frozen foregut cells in hydrogels printed on the pillar plate and dispensed in the 24-well plate was measured. Interestingly, the amount of albumin secreted from HLOs differentiated from the three different conditions was almost identical, with the CV values of 15.8% from 50 μL Matrigel dome in the 24-well plate, 21.5% from 4 μL Matrigel spot on the 36PillarPlate, and 5.4% from 5 μL alginate-Matrigel spot on the 36PillarPlate (**Fig. 5A**). This result clearly demonstrates high reproducibility of HLO function in the mixture of alginate and Matrigel on the pillar plate at 10-fold reduced volume. After generating HLOs on the 36PillarPlate, the expression of a hepatic cell marker, hepatocyte nuclear factor-4α (HNF4α), and an epithelial cell marker, E-cadherin, were detected in HLOs by immunofluorescence staining (**Fig. 5B**). There was no difference in the expression of HNF4α and E-cadherin for HLOs cultured on the pillar (**Fig. 5B** middle rows) as compared to those cultured in a standard Matrigel dome in the 24-well plate (**Fig. 5B** the bottom row). The HLOs consist of multiple hepatic cell types including hepatocyte-, stellate-, and Kupffer-like cells^7,18^. Six iPSC lines from different donors were successfully differentiated into HLOs on the 36PillarPlate and their albumin secretion was measured by an ELISA assay (**Suppl. Fig. 1E**). As a result, all HLOs tested successfully secreted varying levels of albumin, indicting the robustness of the HLO culture protocol. In addition to albumin secretion, micro-anatomical characterization of HLOs has been demonstrated by measuring the intake of bile acid on the 36PillarPlate (**Fig. 5C**). Our HLO model has the capability of bile transportation from the outside and accumulated into the intra-lumen^7,18^. Bile acid excretion is an important detoxification mechanism in the liver since accumulation in hepatocytes can cause the toxic response of cholestasis^22,23^. To monitor the dynamics of bile transport activity in HLOs, HLOs on the 36PillarPlate were treated with cholyl-lysyl-fluorescein (CLF), which is a fluorescein-labeled bile acid with a biological behavior resembling naturally occurring cholylglycine. Time-lapse imaging was performed by incubating the HLOs on the 36PillarPlate with CLF in a 384-well plate for 24 hours. As a result, gradual accumulation of CLF in the lumen of HLO was observed over time, indicating the ability of HLOs to take up bile acid from the outside and excrete it inside HLO.

**Figure 5.**
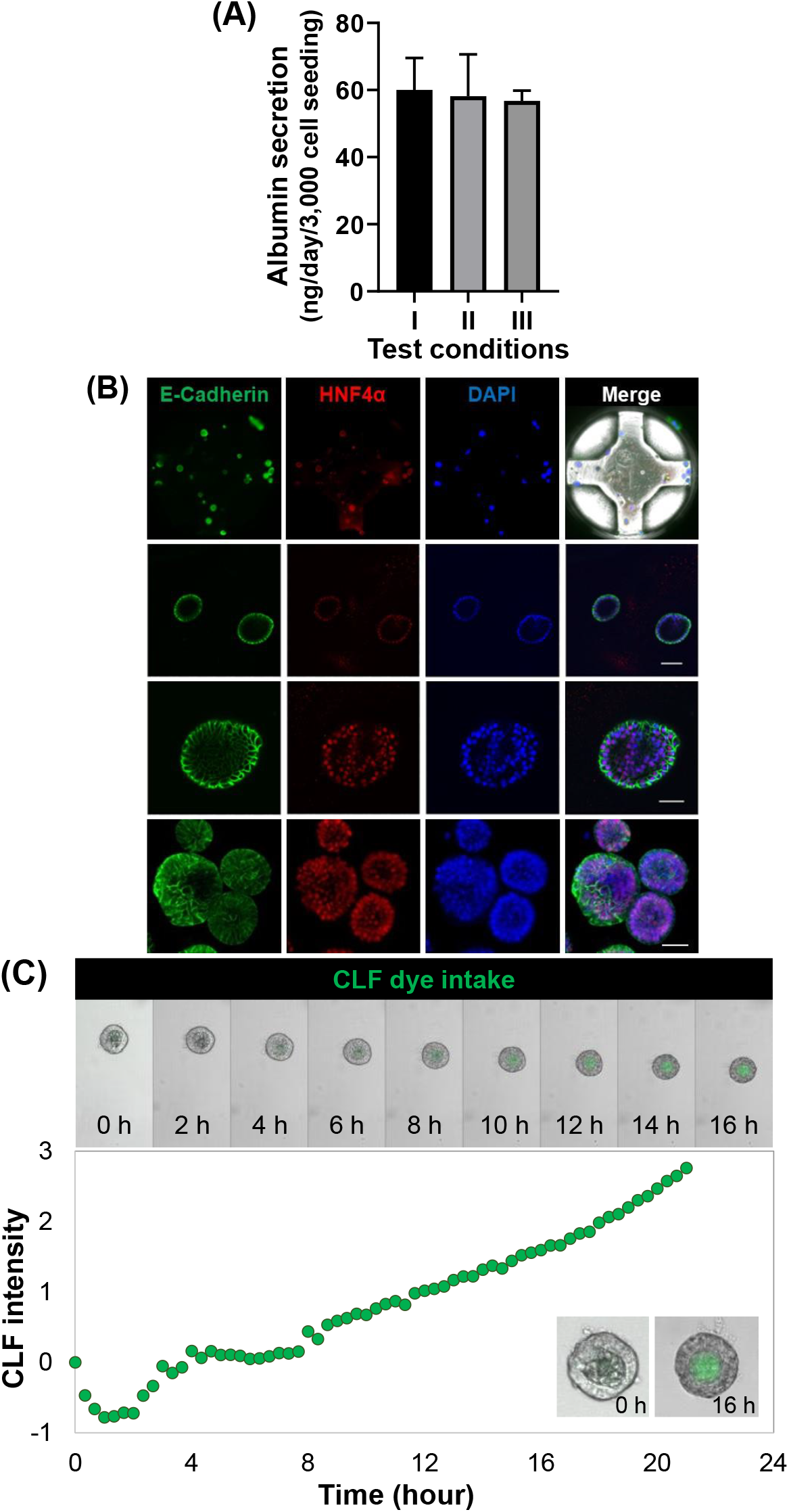
Functionality of static cultured HLOs on the pillar plate. **(A)** Albumin secretion from human liver organoids (HLOs) measured with an albumin human ELISA kit: I) 50 μL Matrigel dome in a 24-well plate, II) 4 μL Matrigel spot on the 36PillarPlate, and III) 5 μL alginate-Matrigel spot on the 36PillarPlate. The same cell number to medium volume ratio was used during HLO differentiation and maturation. The level of albumin secretion from the three test conditions was similar with the CV values of 15.8% from (I), 21.5% from (II), and 5.4% from (III), demonstrating the superior reproducibility of HLO function in the mixture of alginate and Matrigel. (n = 3 pillar plates with 16 pillars/plate) **(B)** Immunofluorescence staining of HLOs generated on the 36PillarPlate and in a 24-well plate. Green, red, and blue colors represent E-cadherin, HNF4α, and nucleus, respectively. Scale bar = 100 μm. The upper row indicates a whole pillar image whereas the middle two rows show low and high magnifications of HLOs on the 36PillarPlate. The bottom row presents an image of HLOs created by standard Matrigel dome culture in the 24-well plate. **(C)** Measurement of bile acid transport function in HLOs on the 36PillarPlate by time-lapse imaging. The green color represents intake of the cholyl-lysyl-fluorescein (CLF) dye in HLOs. The intensity of CLF was quantified by ImageJ.

Incidence of hepatic steatosis, including non-alcoholic fatty liver disease (NAFLD), is increasing, leading to increased morbidity and mortality^7,24,25^. Modeling hepatic steatosis with HLOs on the pillar plate has potential applications in HTS of drug candidates. Using HLOs on the 36PillarPlate, steatohepatitis was successfully simulated by exposing HLOs to fatty acid in the 36PerfusionPlate for 72 hours (**Figs. 6A-C**). Briefly, HLOs on day 18 were dispensed with Matrigel on the 36PillarPlate, which was followed by sandwiching the 36PillarPlate onto the 36PerfusionPlate with 300 μM oleic acid in hepatocyte maturation medium for 72 hours. The capacity of lipid accumulation in HLOs was evaluated by BODIPY-based lipid staining and image analysis (**Fig. 6A**). Further high magnification confocal imaging revealed that fine lipid droplets accumulated in the cytoplasm of HLOs after treatment with oleic acid (**Figs. 6B-C**, P<0.0001, Unpaired t test). Green dots on the pillar represent accumulated lipids in HLOs stained with BODIPY dye whereas purple represents cytoskeleton stained with SiR-actin and blue dots indicate DAPI-stained nuclei.

**Figure 6.**
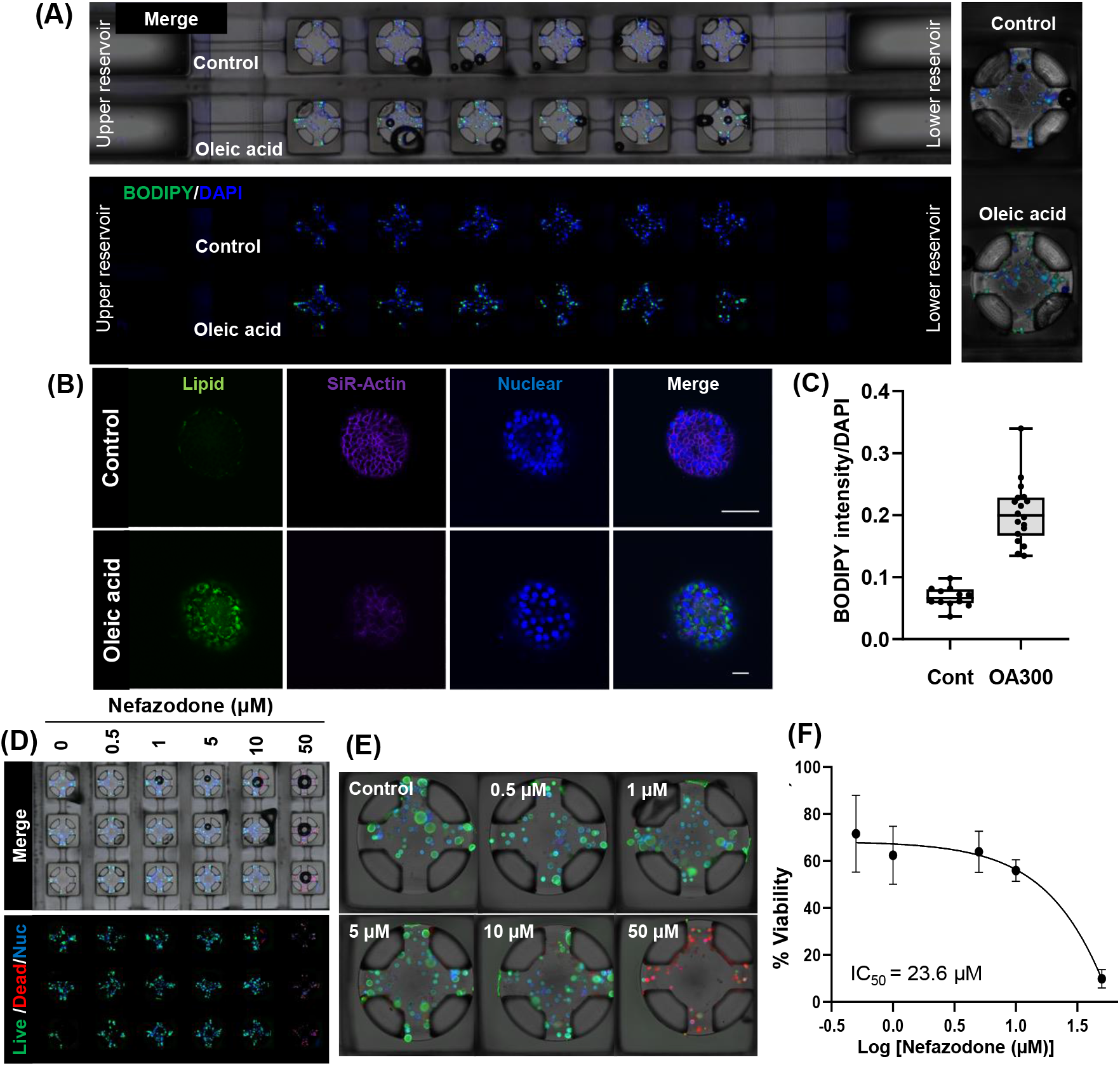
Treatment of static cultured HLOs on the pillar plate with compounds. **(A)** Induction of steatohepatitis in HLO by fatty acid treatment. The top and bottom panel of the pictures indicate HLOs on the 36PillarPlate treated with growth medium (control) and 300 μM sodium oleate (OA300) for 72 hours in the 36PerfusionPlate and imaged by bright-field and fluorescence microscopy. The left panel is blow-up images of the representative pillar with HLOs. Green dots represent accumulated lipids stained with BODIPY dye whereas blue dots indicate DAPI-stained nuclei. **(B)** Confocal imaging of lipid accumulation on the 36PillarPlate. The upper row represents control HLO whereas the bottom row represents HLO treated with 300 μM sodium oleate for 72 hours. Green, purple, and blue colors represent lipid, SiR-actin, and nucleus staining, respectively. Scale bar = 100 μm. **(C)** Quantification of lipid accumulation. The intensity of BODIPY was quantified by ImageJ and normalized by DAPI intensity. **(D, E)** Measurement of hepatotoxicity with HLOs on the pillar plate: (D) low magnification image of the whole pillar plate and (E) high magnification image of each pillar. HLOs on the 36PillarPlate were treated with 0 – 50 μM of nefazodone in the 36PerfusionPlate for 72 hours. Live/dead staining of HLOs was performed with calcein AM (green) and ethidium homodimer-1 (red). Green, red, and blue colors represent live cells, dead cells, and nuclei, respectively. **(F)** Measurement of dose response of nefazodone. After nefazodone treatment for 72 hours, the viability of HLOs was determined by CellTiter-Glo^®^ luminescent cell viability assay kit.

For the assessment of drug-induced liver injury (DILI)^18^, the 36PillarPlate with HLOs was sandwiched onto the 36PerfusionPlate containing nefazodone and incubated for 72 hours (**Figs. 6D-F**). Nefazodone, an antidepressant, with human C_max_ value of 1.34 μM and rat oral LD_50_ value of 980 mg/kg, has been known to cause idiosyncratic DILI^26,27^. After drug exposure, the viability of HLOs was measured by detaching the 36PillarPlate and sandwiching it onto a new 384-deep well plate containing calcein AM and ethidium homodimer-1 (**Fig. 6E**). Green and red dots on the pillar represent live and dead cells, respectively. Cell viability decreased in response to increase in nefazodone concentrations. The dose response curve and the IC_50_ value of nefazodone were determined using CellTiter-Glo^®^ luminescent cell viability assay kit (**Fig. 6F**). Similar to the results of image analysis, the cell viability decreased in a nefazodone dose-dependent manner. The IC_50_ value of nefazodone obtained from the 36PillarPlate (six replicates per dosage) was 23.6 μM, which was comparable to the IC_50_ of 29.4 μM obtained for 3D human liver microtissue^28^.

### Cell spheroid transferring to the pillar plate for human intestine organoid (HIO) culture

Unlike HLOs, generation of certain organoids such as intestine and brain organoids require direct cell-to-cell contact by creating cell spheroids prior to embedding in biomimetic hydrogels^8,29^. These cell spheroids are difficult to dispense robustly using conventional liquid handling machines due to rapid settling down of spheroids in aqueous solutions. In addition, most liquid handling machines are incapable of uniformly dispensing cells in hydrogels, particularly in temperature-sensitive hydrogels such as Matrigel, due to lack of a chilling dispensing head. To address this issue, we developed a new method of transferring single spheroids from an ultralow attachment (ULA) 384-well plate to the pillar plate by simple sandwiching and inverting for highly uniform differentiation of organoids (**Suppl. Fig. 2B**)^30^. Briefly, PSCs were differentiated into monolayers of mid-hindgut cells, which were then disassociated into a single cell suspension. Approximately 3,000 cells per well were dispensed into a Nexcelom ULA 384-well plate, resulting in formation of spheroids (200 μm in diameter). Spheroids were transferred *en masse* directly from the ULA wells onto the pillars of a 36PillarPlate. Mid-hindgut aggregates were differentiated into HIOs over the course of 40 days (**Fig. 7A**). The overall success rate of spheroid transfer to the pillar plate was between 95 - 100%. This approach is amenable to HTS and eliminates the labor-intensive spheroid pipetting step. After HIO differentiation and maturation for 40 days on the pillar plate, organoid clearing and staining protocols were established for *in situ* organoid imaging on the pillar plate and demonstrated intestinal epithelium development as well as cell-cell tight junction formation in HIOs (**Figs. 7B and 7C**).

**Figure 7.**
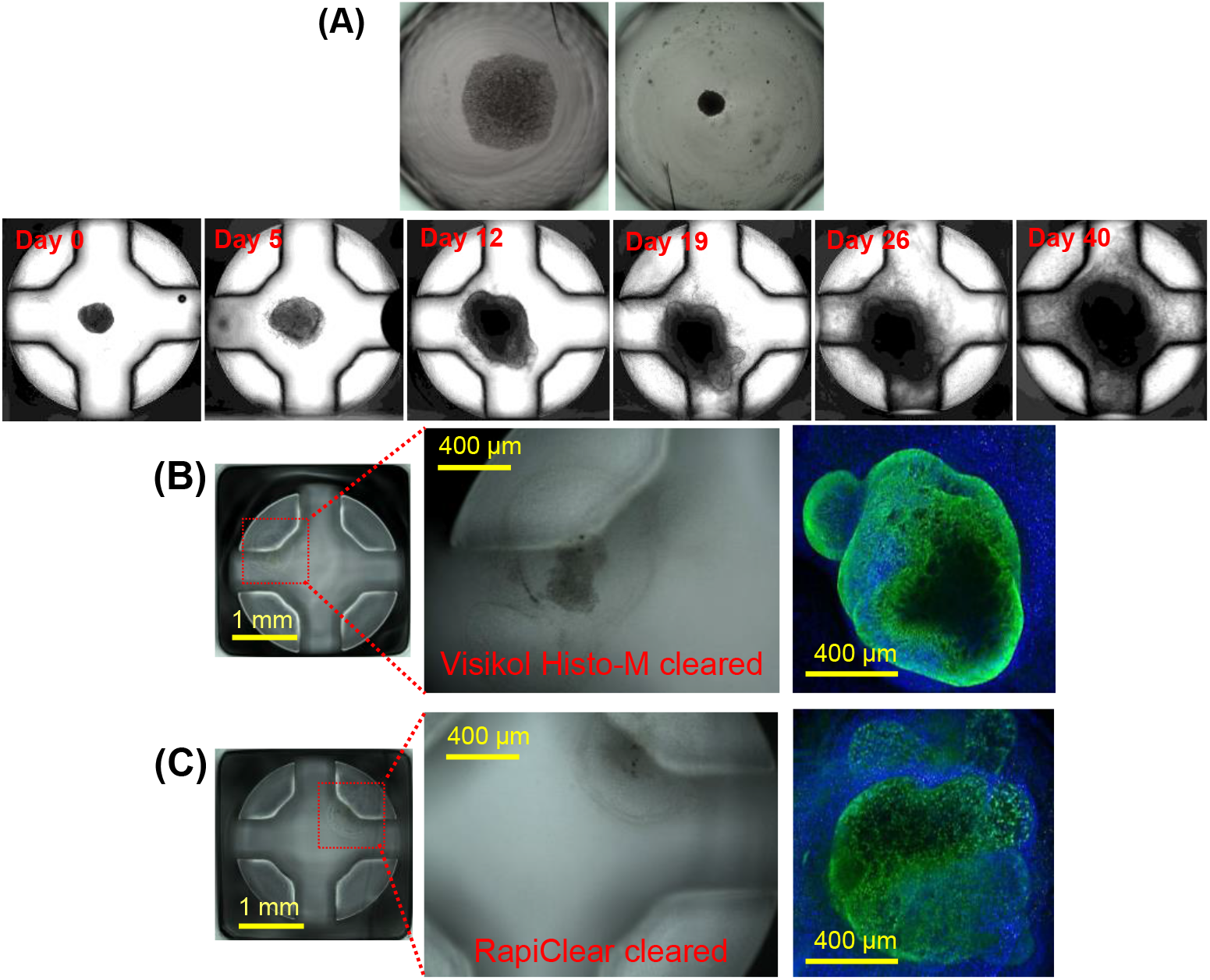
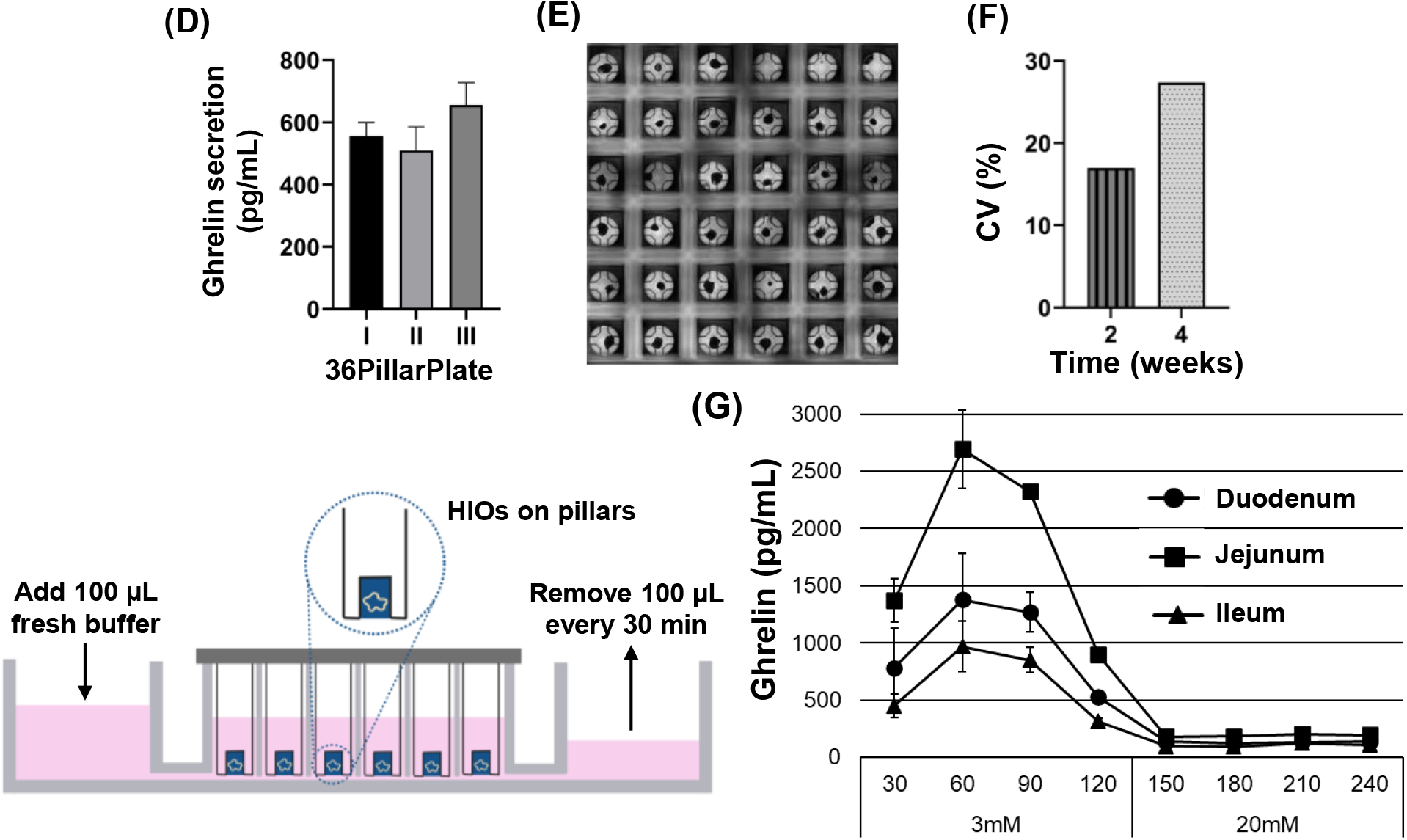
Uniform differentiation of human intestine organoids (HIOs) on the pillar plate by transferring single mid-hindgut cell spheroids from an ultralow attachment (ULA) 384-well plate to the 384PillarPlate. **(A)** Bright-field images of frozen mid-hindgut cell spheroids cultured in Nexcelom ULA 384-well plate for 2 days, transferred to the 384PillarPlate, and differentiated into HIOs. Single spheroid in diameter of 200 μm was robustly transferred from the ULA 384-well plate to the 384PillarPlate (95 - 100% success rate achieved reproducibly). Transferred spheroids were all viable and uniform. **(B)** HIOs generated from frozen mid-hindgut cell spheroids in Nexcelom ULA 384-well plate transferred and encapsulated in 2-fold diluted Matrigel on the 384PillarPlate and cultured in gut differentiation medium for 40 days (4x and 10x magnification). Z-stacked fluorescent images were obtained after staining with E-cadherin (green) for cell-cell tight junction formation and DAPI (blue) for nucleus (10x). **(C)** HIOs stained with CDX2 (green) for intestinal epithelium development and DAPI (blue) for nucleus (10x). Images captured by Nikon A1 Rsi confocal microscope after Visikol Histo-M or RapiClear clearing. **(D)** Amounts of ghrelin secreted from HIOs on the 36PillarPlate after incubation in normal media and measured by a human Ghrelin ELISA kit (ThermoFisher). (n = 6) Triplicate tests were performed to measure ghrelin secretion. **(E)** Stitched image of HIOs on the 36PillarPlate after 14 days of culture. **(F)** Measurement of coefficient of variation (CV) with cell viability to determine reproducibility of HIO generation. **(G)** Dynamic Ghrelin secretion from HIOs on the 36PillarPlate observed using the 36PerfusionPlate with manual addition and sampling of nutrient buffer. Ghrelin secretion was elevated during low glucose (3 mM) conditions, and abruptly ceased at high glucose (20 mM) concentrations.

To test HIOs for functionality we determined if organoids were capable of secreting the satiety hormone, ghrelin. We first tested for ghrelin secretion from HIOs that we manipulated to contain high numbers of hormone-secreting enteroendocrine cells as previously described^31^. In static culture, HIOs cultured in normal media secreted very consistent, low levels of ghrelin (**Fig. 7D**). After 4-week culture of HIOs on the 36PillarPlate, 8 - 14 % of HIOs on the pillar plate were detached due to degradation of Matrigel by proteases secreted whereas 5 - 11 % of mid-hindgut aggregates were not differentiated into HIOs properly potentially due to the small number of cells (**Fig. 7E**). Nonetheless, uniform functionality of HIOs on the pillar plate was achieved repeatedly with CV values below 25% (**Figs. 7D and 7F**). We then determined if the perfusion plate could be used to dynamically study the function and physiologic responses of HIOs *in vitro*. Ghrelin is a hormone that regulates feeding behavior in humans. Under fed conditions, ghrelin levels are low, but they increase under fasting conditions when circulating nutrient levels drop. To investigate if HIOs respond to nutrient levels like the human intestine, we used the 36PillarPlate/36PerfusionPlate system to dynamically control the glucose levels on the pillars from 3 mM (fasting) to 20 mM (fed) (**Fig. 7G**). HIOs were cultured in 5 mM glucose containing media overnight (equilibration media) and then shifted to fasting media by removing the equilibration media from upper and lower reservoirs (UR and LR) and adding 500 μL of fasting media (3 mM glucose) to the UR. For each 30-minute timepoint, 100 μL of the fresh low glucose buffer solution was added in the URs and the same volume of solution was removed from the LRs. We observed a dynamic increase in ghrelin secretion in response to fasting conditions. To determine if HIOs would respond to feeding by shutting down ghrelin secretion, we replaced the fasting media with identical media containing 20 mM glucose. This resulted in a precipitous drop in ghrelin secretion. Interestingly, jejunum HIOs responded to glucose more significantly as compared to duodenum and ileum HIOs. These results demonstrate that we can generate uniform organoids scalably by transferring spheroids to the pillar plate and monitor organoids either by *in situ* imaging, or in real-time using a simple fluidic modality.

## Discussion

The vast majority of organoids have been cultured in static conditions with PSCs encapsulated in biomimetic hydrogels such as Matrigel^1,2^. Single PSCs suspended in Matrigel or PSC spheroids mixed with Matrigel are loaded manually with a multichannel pipette in 6-/24-well plates for differentiation and maturation of PSCs into organoids. Once the diameter of organoids reaches around 1 mm, organoids can be cultured in petri dishes and spinner flasks with shaking to minimize cell death in the core due to diffusion limitations of oxygen and nutrients. A primary concern in the field is variability of organoids within the same isolation and from donor-to-donor, and in the case of PSC-derived organoids, within the same differentiation and between separate differentiations^19,32^. This can be due to patient-to-patient genetic variability and differences in laboratory techniques despite the usage of identical reagents and cell lines. A simple but resource-heavy approach to mitigate these issues is significant culture scale-up combined with automated analysis of only organoids that pass certain criteria^33^. Variability can also be reduced by inserting selection steps during the organoid formation process^34^, or by incorporating mechanical and fluidic handling^35^. We addressed the variability issue by introducing robotic cell printing and miniaturization of organoid culture in static and dynamic culture on the pillar/perfusion plate platform and achieved overall CV values below 25%.

Although there are several 3D cell culture systems developed^36,37^, these are not amenable to organoid culture and analysis for high-throughput, predictive compound screening. For example, ULA well plates and hanging droplet plates marketed by Corning and InSphero have been used for relatively short-term culture of cell spheroids, which is different from long-term cultured organoids in hydrogels. Almost all organoid cultures require encapsulation of PSCs in biomimetic hydrogels for maturation, and high-density well plates are unsuitable for organoid culture in hydrogels due to difficulty in changing growth media. Transwell inserts from Corning are designed primarily for layered cell co-culture, which are low throughput to seed cells in layers and cumbersome to change growth media. Recent advances in “3D bioprinting” from Organovo and others offer new opportunities for creating highly organized multicellular tissue constructs *in vitro* mainly by using micro-extrusion robots to precisely dispense multiple cell types in natural and synthetic hydrogels (“bioinks”) layer-by-layer^38^. However, photopolymerization often used for 3D bioprinting may be toxic to PSC-derived organoids due to oxygen radicals generated from photoinitiators^39^. Current extrusion-based 3D bioprinting approaches need to be miniaturized for organoid culture to save expensive human cell sources, to reduce cell death in the core of bioprinted tissues due to limited diffusion of oxygen and nutrients, and to improve throughput of organoid imaging *in situ*. Robotic cell spheroid-based 3D bioprinting (“Kenzan” technology) marketed by Cyfuse could be used to precisely position mature organoids isolated from Matrigel in microtiter well plates^40^. However, it would take a long time to pick up a single spheroid at a time and dispense it individually in a microtiter well plate with Matrigel for HTS assays. MPSs with microfluidic channels and chambers from Mimetas, Nortis, and Emulate have been developed and widely studied to recapitulate micro-vessel structures *in vivo*^41^. These MPS systems contain cultured 3D cells in microfluidic chambers representing an organ and microfluidic channels providing in- and out-flow for growth media, thus leading to better retention of *in vivo-* like cell functions as compared to static cultures. In addition, MPSs have been developed to model adsorption, distribution, metabolism, elimination, and potential toxicity (ADMET) of chemicals *in vitro*,^42,43^ but can also be used to control cell differentiation as well as nutrient exposure. The effects of xenobiotics on developing organoids^44^ as well as nutrient conditions^45^ have been modeled under highly controlled environments enabled by MPSs. The physical stress of fluid flow itself has been shown to impart differing effects on organoids such as enhanced nutrient exposure/waste removal^45^, formation of endothelial cells from nascent mesenchyme^11^, and neurite outgrowth^46^. However, many examples of MPSs suffer from inherent low throughput and poor maneuverability for organoid culture due to tedious manual steps necessary for cell loading and media change, cumbersome pump systems required to connect different organ modules, or required highly specialized skills and the purchase of expensive proprietary equipment^19,41,47^. These issues represent a major gap in MPS development for organoid research. Thus, the design of microfluidic channels and chambers may need to be much simpler to provide higher throughput in organoid culture, easier cell loading and media change, and higher compatibility with existing HTS equipment.

To the best of our knowledge, there is no 3D HTS system existing for dynamic organoid culture and analysis. Current organoid cultures heavily rely on 6-/24-well plates, petri dishes, and spinner flasks, which are labor-intensive and difficult to automate for HTS. Unlike other 3D cell culture systems, our pillar/perfusion plate platform allows for combining rapid 3D bioprinting with “microfluidic-like” features for static and dynamic organoid cultures. There are several unique features of the pillar/perfusion plates for HTS, which include high-throughput, high reproducibility, and cost-effective cell printing protocols that can be used for controlling cellular microenvironments for disease modeling. Highly reproducible, high-throughput precision printing allows for testing a variety of organoid culture conditions with individual and mixtures of compounds, which makes it well suited for early-stage HTS of compound libraries. The pillar/perfusion plate platform requires relatively small amounts of cells, hydrogels, ECMs, growth factors, compounds, and reagents for creating and evaluating organoids. Multiple organoids with physiologically relevant characteristics of the tissue of origin can be created on a single pillar plate by static and dynamic culture that could provide predictive insight into potential organ-specific toxicity of compounds. The pillar/perfusion plate is manufactured by injection molding with polystyrene, which is nontoxic and presents no concerns for nonspecific compound adsorption. In addition, it is optically clear for direct visualization of organoids on the pillars for *in situ* high-content cell staining and imaging. Cell image acquisition from bioprinted organoids is easy and straightforward because the whole sample depth fits within the focus depth of a normal objective (4x and 10x magnification). In addition, the pillar/perfusion plate is highly flexible for biological assays with organoids. For example, organoids on the 36PillarPlate can be cultured in the 384DeepWellPlate or the 36PerfusionPlate to simulate static or dynamic conditions. Unlike traditional MPSs, the 36PillarPlate with multiple organoids can be easily detached from the 384DeepWellPlate or the 36PerfusionPlate and then sandwiched onto conventional 384-well plates containing cell-staining reagents for high-throughput assays. The perfusion plate requires no pumps or tubes, which makes it easy to change growth media for long-term organoid culture as well as to examine compound-organoid and organoid-organoid interactions. Multiple organoid types can be created on the 36PillarPlate and then connected using the 36PerfusionPlate to simulate complex diseases with multiple organoids involved. Moreover, the pillar/perfusion plate built on the footprint of standard 384-well plates is compatible with existing HTS equipment such as fully automated fluorescence microscopes and microtiter well plate readers, which is an important feature for developing HTS assays with organoids.

### Outlook

We have successfully manufactured the pillar/perfusion plate *via* injection molding of polystyrene and demonstrated static and dynamic HLO and HIO culture with functional assays. The pillar/perfusion plate maintained long-term organoid culture with bioprinted single cells suspended in the mixture of alginate and Matrigel and cell spheroids in Matrigel by supporting either static culture with growth media in the deep well plate or dynamic culture with a flow of growth media through the perfusion well plate without the use of pumps or tubes. Developed cell printing and encapsulation protocols were highly flexible and allowed for culturing multiple organoids in Matrigel on the pillar plate, consequently providing more insight into potential organ-specific toxicity of compounds. Our microarray 3D bioprinting technology demonstrated on the pillar/perfusion plate represents a unique strategy of printing PSCs and organoids in biomimetic hydrogels on the pillar plate rapidly. Human organoids on the pillar/perfusion plate platform could recapitulate tissue development and maintain high tissue functions by mimicking *in vivo* microenvironments. In addition, tissue functions and mechanistic actions of compounds could be replicated and elucidated *in vitro* through high-throughput, high-content organoid analysis. Bioprinted organoids were combined with high-content, whole organoid imaging to better understand functions of organoids generated and cytotoxicity of compounds. Thus, our microarray 3D bioprinting technology could address the unmet need in organoid research by combining rapid printing of PSCs on the pillar plate, differentiating and maturing into organoids in static and dynamic culture to mimic the physiological microenvironment of tissues inside the human body while enhancing throughput and maneuverability dramatically for predictive screening of compounds. We envision that bioprinted human organoids in the pillar/perfusion plate could provide highly predictive toxicity and efficacy information needed in preclinical evaluations of compounds or prioritize environmental toxicants. Thus, our unique approach could offer wide industrial adoption of organoid-based assays for HTS.

## Methods

### Fabrication of pillar/perfusion plates

The 384PillarPlate, the 384DeepWellPlate, the 36PillarPlate, and the 36PerfusionPlate (**Figs. 2 & 3**) were manufactured by injection molding of polystyrene and functionalized with proprietary polymer solutions (Bioprinting Laboratories Inc., Denton, TX, USA). The 384PilllarPlate and the 36PillarPlate contain 384 pillars and 36 pillars (4.5 mm pillar-to-pillar distance, 11.6 mm pillar height, and 2.5 mm outer and 1.5 mm inner diameter of pillars), respectively. The 384DeepWellPlate and the 36PerfusionPlate have 384 complementary wells (3.5, 3.5, and 14.7 mm well width, length and depth, and 4.5 mm well-to-well distance) and 36 wells (3.4, 3.4, and 11.9 mm well width, length and depth, and 4.5 mm well-to-well distance) with microchannels and reservoirs, respectively.

### Simulation of flow patterns within the perfusion plate using SolidWorks

To ensure uniform mixing and avoid diffusion limitation on the pillars inserted in the perfusion wells, a 3D computational model for the flow rate and profile has been developed using the SolidWorks software package (SolidWorks Research Standard and Simulation Premium 2022, Waltham, MA, USA). To estimate the fluid dynamics within the 36PerfusionPlate, the transport phenomenon was simulated using a simplified incompressible Navier-Stokes equation below:

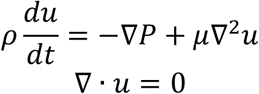

where, ρ is water density [ML^−3^], u is flow velocity [LT^−1^], P is pressure in the UR [ML^−1^T^−2^], and μ is dynamic viscosity of water [ML^−1^T^−1^]. The geometry of the 36PerfusionPlate was simplified to model only six rectangular perfusion wells with two reservoirs in a single row connected by microchannels (**Fig. 3F**). The microchannels have a dimension of 0.4 mm × 0.4 mm. The model assumed that the 36PillarPlate is sandwiched onto the 36PerfusionPlate. There is a 0.5 mm gap from the bottom of perfusion wells to the top of pillars. In addition, it was assumed that the water-air boundary at the top of the perfusion wells acted as a wall. By changing the level of water in the UR and the LR (i.e., inlet and outlet pressure) on a digital rocker, we determined the velocity profiles within the perfusion wells to avoid diffusion limitation of growth media on the 36 pillars.

### Bioprinted foregut cell culture in the mixture of alginate and Matrigel on the pillar plate to create human liver organoids (HLOs)

HLOs were generated on the 36PillarPlate with modified protocols previously described (**Suppl. Fig. 2A**)^7,18^. Briefly, definitive endoderm was generated on day 3 by differentiating multiple iPSCs (Cincinnati Children’s Hospital Medical Center) using 50 ng/mL bone morphogenetic protein 4 (BMP4) and 100 ng/mL Activin A along with 0.2% fetal bovine serum (FBS) on day 2^7,18^. This was followed by inducing foregut cells using 500 ng/mL fibroblast growth factor 4 (FGF4) and 3 μM CHIR99021 (GSK3 inhibitor). Foregut cells were collected with mesenchymal cells by gentle pipetting and cryopreserved for long-term storage. Frozen foregut cells were thawed, centrifuged to remove cryoprotectants, and mixed with the mixture of cold alginate and Matrigel™ matrix (Corning Inc., NY, USA) on ice to achieve a final concentration of 600,000 cells/mL in 0.75% alginate and 6-8 mg/mL Matrigel. The suspension of foregut cells in alginate-Matrigel mixture was printed on top of the 36PillarPlate at a volume of 5 μL (3,000 cells per pillar) using a 3D bioprinter (ASFA™ spotter from MBD Korea) while maintaining the slide deck at 4°C to prevent water evaporation during printing. For hydrogel gelation, the 36PillarPlate was sandwiched with an 384DeepWellPlate containing 10 mM CaCl_2_ and incubated in a 5% CO_2_ incubator at 37°C for 10 - 15 minutes. For HLO formation, the 36PillarPlate was sandwiched onto a 384DeepWellPlate containing 80 μL of organoid formation medium in advanced DMEM/F12 with 2% B27, 1% N2, 10 mM HEPES, 1% Glutamax or L-glutamine, 1% Pen/Strep, 5 ng/mL fibroblast growth factor 2 (FGF2), 10 ng/mL vascular endothelial growth factor (VEGF), 20 ng/mL epidermal growth factor (EGF), 3 μM CHIR99021, 0.5 μM A83-01, 50 μg/mL ascorbic acid, and the CEPT cocktail^21^, and incubated in the CO_2_ incubator for 4 days with medium change every other day. For HLO differentiation, the 36PillarPlate was separated and sandwiched onto another 384DeepWellPlate containing 80 μL of hepatic canaliculi formation medium in advanced DMEM/F12 with 2% B27, 1% N2, 10 mM HEPES, 1% Glutamax or L-glutamine, 1% penicillin/streptomycin, and 2 μM retinoic acid (RA), and incubated in the CO_2_ incubator for 4 days with medium change every other day. To promote the polarization of organoids and increase size and complexity of the bile canaliculi and the pericanalicular sheaths, the encapsulated cells were incubated with RA^48^. After RA treatment, HLOs were cultured in hepatocyte maturation medium in hepatocyte culture medium (Lonza, catalog no. CC-3198) with 10 ng/mL hepatocyte growth factor (HGF), 0.1 μM dexamethasone (DEX), and 20 ng/mL oncostatin M (OSM) for 10 days.

### Mid-hindgut cell spheroid culture in Matrigel on the pillar plate to create human intestine organoids (HIOs)

HIOs were generated on the 384PillarPlate with modified protocols previously described (**Suppl. Fig. 2B**)^49^. Briefly, iPSCs were maintained in mTeSR (StemCell Technologies) in monolayer and passaged every 7 days using Gentle Cell Dissociation Reagent (StemCell). iPSCs were treated with Accutase (StemCell) and plated on to 24-well plates coated with ESC-qualified Matrigel (Corning) in mTeSR supplemented with Y-27632 (Vendor). iPSCs were then differentiated into definitive endoderm by treatment with 100 ng/mL Activin A (R&D) and 15 ng/mL BMP4 (R&D) in RPMI supplemented with nonessential amino acids (NEAA, Gibco) for 1 day, followed by 100 ng/mL Activin A in RPMI with NEAA and 0.2% FBS (Gibco) for 1 day, and 100 ng/mL Activin A in RPMI with NEAA and 2% FBS for 1 day. The resulting definitive endoderm was subjected to 3 μM CHIR 99021 and 500 ng/mL FGF4 (R&D) in RPMI with NEAA and 2% FBS for 4 days. The resulting mid-hindgut cell monolayer was dissociated using Accutase for 8 - 10 minutes. The resulting single cell suspension was cryopreserved in Cell Banker 1 (Amsbio) for long-term storage. Frozen mid-hindgut cells were thawed, centrifuged to remove cryoprotectants, and resuspended in Gut maturation medium (consisting of Advanced DMEM/F12 with N2, B27, 15 mM HEPES, 2 mM L-glutamine, 100 UI/mL penicillin/streptomycin) supplemented with 100 ng/mL EGF (R&D) and the CEPT cocktail^21^. The single mid-hindgut cell suspension was aggregated in a ULA 384-well plate (Nexcelom) for 4 days with each spheroid consisting of approximately 3,000 cells. For HIO culture on the pillar plate, 2-fold diluted, cold Matrigel was printed on the 36PillarPlate (5 μL per pillar) using the 3D bioprinter and stored at room temperature for 5 minutes for partial gelation. For spheroid transfer, the 36PillarPlate with Matrigel was sandwiched onto the ULA 384-well plate with cell spheroids, and then the sandwiched plates were inverted with the pillar plate down. After incubating the inverted, sandwiched plates for 20 minutes in the CO_2_ incubator for complete Matrigel gelation, the 36PillarPlate with cell spheroids encapsulated in Matrigel was separated and sandwiched onto a 384DeepWellPlate containing 80 μL of Gut maturation medium supplemented with 100 ng/mL EGF per well. HIOs were generated by culturing the cells in a humidified CO_2_ incubator for 4 - 5 weeks with medium changes every other day.

### Operation of the perfusion plate for dynamic cell culture

For dynamic cell culture in the 36PerfusionPlate, 450 μL of fresh growth medium was added in each UR and LR (**Suppl. Fig. 3E**). After covering the 36PerfusionPlate with a lid to prevent water evaporation and contamination, the 36PerfusionPlate was incubated in the CO_2_ incubator for 10 minutes until the level of the growth medium in the perfusion wells reached equilibrium. The 36PillarPlate with cells encapsulated in hydrogels was sandwiched onto the 36PerfusionPlate, and then the sandwiched plates were placed on a digital rocker (OrganoFlow^®^ L, Mimetas) in the CO_2_ incubator. The tilt angle and frequency of the rocker are set at 10° and 1 minute. After 1 - 2 days of cell culture, old growth medium was removed from each LR at 10° tilt angle, and then 900 μL of fresh growth medium was added in each UR immediately. This simple medium change can be repeated for long-term organoid culture.

### Measurement of cell viability using CellTiter-Glo^®^ luminescent cell viability assay kit

The 36PillarPlate with HLOs was sandwiched with an opaque white 384-well plate containing 20 μL of CellTiter-Glo^®^ Reagent from CellTiter-Glo^®^ cell viability kit (Promega, Madison, WI, USA) and 20 μL of cell culture medium in each well to measure cellular adenosine triphosphate (ATP) levels. To induce cell lysis, the sandwiched pillar/well plates were placed on an orbital shaker for 25 minutes. After stabilizing the luminescence for 15 minutes at room temperature, the luminescence signals were recorded using the microtiter well plate reader (Synergy H1) at an emission wavelength of 560 nm. A standard calibration curve was generated using ATP disodium salt (Sigma Aldrich) at a concentration range of 10 nM to 1 μM.

### Measurement of cell viability using calcein AM and ethidium homodimer-1

The HLOs on the 36PillarPlate were rinsed once with a saline solution and then stained with 80 μL of a mixture of 2 μM calcein AM, 4 μM ethidium homodimer-1, and one drop of NucBlue^®^ Live ReadyProbes^®^ Reagent (ThermoFisher Scientific) in a 384DeepWellPlate for 1 hour at room temperature. After staining, the 36PillarPlate was rinsed twice with the saline solution, and fluorescent images were acquired in high throughput with Keyence BZ-X710 automated fluorescence microscope (Keyence, Osaka, Japan).

### Measurement of albumin secretion from HLOs

To measure the level of albumin secretion from HLOs on pillars, 80 μL of culture medium in the 384DeepWellPlate was collected after 48 hours of incubation with HLOs encapsulated in alginate-Matrigel mixture on the pillar plate. The culture medium was centrifuged at 250 g for 3 minutes to remove any debris, and the resulting supernatant was assayed with human albumin ELISA kit (EHLAB, ThermoFisher Scientific) according to the manufacturer’s instruction. To quantify the number of HLOs on each pillar and normalize the amount of albumin secreted by the HLO number, images were captured with the Keyence microscope.

### Immunofluorescence staining of whole HLOs on the pillar plate

HLOs on the 36PillarPlate were rinsed by sandwiching the 36PillarPlate onto a 384DeepWellPlate with 80 μL of 1x phosphate-buffered saline (PBS) at room temperature for 15 - 20 minutes on a slow-speed orbital shaker. The HLOs were fixed with 80 μL of 4% paraformaldehyde (PFA) solution in the 384DeepWellPlate for 30 - 60 minutes at room temperature. The fixed HLOs were washed with 0.1% (w/v) sodium borohydride in PBS for 15 minutes to reduce background due to free aldehyde. After washing the 36PillarPlate with 80 μL of 1x PBS in the 384DeepWellPlate at room temperature twice for 15 - 20 minutes each on the slow-speed shaker, HLOs were permeabilized with 80 μL of 1% Triton X-100 in PBS for 15 minutes followed by 0.1% Tween 20 for 15 minutes in the 384DeepWellPlate at room temperature on the slow-speed shaker. After washing the 36PillarPlate once with 80 μL of 1x PBS with 0.5% Triton X-100 in the 384DeepWellPlate at room temperature for 15 - 20 minutes on the slow-speed shaker, HLOs were exposed to 80 μL of 2% normal donkey serum (NDS) diluted in 1x PBS with 0.5% Triton X-100 (blocking buffer) for overnight at 4°C or 2 - 4 hours at room temperature to prevent non-specific binding. For primary antibody staining, HLOs were treated with 80 μL of 5 - 10 μg/mL diluted primary antibody in the blocking buffer for 1 day at 4°C on the slow-speed shaker. To remove unbound primary antibody, HLOs were rinsed three times with 80 μL of 1x PBS with 0.5% Triton X-100 for 15 - 20 minutes each at room temperature on the slow-speed shaker. For secondary antibody staining, HLOs were exposed to 80 μL of 200x-diluted, fluorophore-conjugated secondary antibody in the blocking buffer for 1 day at 4°C or 2 hours at room temperature on the slow-speed shaker. To remove unbound secondary antibody and stain with DAPI, HLOs were stained with 80 μL of 1 μg/mL DAPI in 1x PBS with 0.5% Triton X-100 for 25 minutes followed by washing with 80 μL of 1x PBS for 15 - 20 minutes at room temperature on the slow-speed shaker. HLOs were cleared by incubating in 30 μL of 1x pre-warmed (at 37°C) RapiClear 1.52 (Sunjin Lab, Taiwan) or Visikol^®^ Histo-M™ (SKU: HM-30, Visikol) in a clear flat-bottom, 384-well plate at room temperature for 0.5 - 1 hour. The stained and cleared HLOs were inspected under a Nikon A1 Rsi confocal microscope.

### Measurement of bile acid transport function in HLOs on the pillar plate by time-lapse imaging

To analyze the function of bile acid transport in HLO, HLOs on the 36PillarPlate were treated with 5 nM cholyl-lysyl-fluorescein (AAT Bioquest Inc., CA, USA) in a 384-well plate. Time-lapse images were captured in 20-minute intervals for 16 hours using Celldiscoverer 7 (Carl Zeiss, Oberkochen, Germany) equipped with 10× objective lens at 37°C and 5% CO_2_. Time-lapse images obtained from the 36PillarPlate were batch-processed using ImageJ (NIH) to extract fluorescence intensity from the entire HLOs.

### Measurement of lipid accumulation in HLOs using BODIPY

Accumulation of lipid in HLOs on the pillar plate was measured using BODIPY^®^ 493/503 (ThermoFisher Scientific). Briefly, HLOs on the 36PillarPlate were treated with the hepatocyte maturation medium in the presence and absence of 300 μM sodium oleic acid (OA300) for 72 hours. Following incubation in the 36PerfusionPlate, HLOs were rinsed three times with warm PBS to remove any residual oleic acid on the cell surface. Lipids accumulated in HLOs and cytoskeleton were stained with 2 μM BODIPY^®^ 493/503 and 1 μM SiR-Actin (Cytoskeleton Inc., CO, USA), respectively. Nuclei were stained with NucBlue^®^ Live ReadyProbes^®^ Reagent. After staining, HLOs on the 36PillarPlate were visualized and scanned using a Nikon A1 inverted confocal microscope (Japan) equipped with 10× objective lens and Keyence BZ-X710 automated fluorescence microscope. The lipid droplet volume was calculated by using ImageJ and normalized with each nucleus signal. Statistical difference in lipid accumulation between control and lipid exposure conditions was determined by Student’s t-test. Statistically significant difference between control and test conditions was indicated by ^*^ for P < 0.05, ^**^ for P < 0.01, and ^***^ for P < 0.001.

### Immunofluorescence staining of whole HIOs on the pillar plate

HIOs on the 36PillarPlate were rinsed by sandwiching the 36PillarPlate onto a 384DeepWellPlate with 80 μL of 1x PBS at room temperature for 15 - 20 minutes on a slow-speed orbital shaker. The HIOs were fixed with 80 μL of 4% paraformaldehyde (PFA) solution in the 384DeepWellPlate for 1 day at 4°C. The fixed HIOs were washed with 0.1% (w/v) sodium borohydride in PBS for 15 minutes to reduce background due to free aldehyde. After washing the 36PillarPlate with 80 μL of 1x PBS in the 384DeepWellPlate at room temperature twice for 15 - 20 minutes each on the slow-speed shaker, HIOs were permeabilized with 80 μL of 1% Triton X-100 in PBS for 2 - 3 hours followed by 0.1% Tween 20 for 30 minutes in the 384DeepWellPlate at room temperature on the slow-speed shaker. After washing the 36PillarPlate once with 80 μL of 1x PBS with 0.5% Triton X-100 in the 384DeepWellPlate at room temperature for 15 - 20 minutes on the slow-speed shaker, HIOs were exposed to 80 μL of 2% normal donkey serum (NDS) diluted in 1x PBS with 0.5% Triton X-100 (blocking buffer) for overnight at 4°C or 2 - 4 hours at room temperature to prevent non-specific binding. For primary antibody staining, HIOs were treated with 80 μL of 5 - 10 μg/mL diluted primary antibody in the blocking buffer for 1 day at 4°C on the slow-speed shaker. To remove unbound primary antibody, HIOs were rinsed three times with 80 μL of 1x PBS with 0.5% Triton X-100 for 15 - 20 minutes each at room temperature on the slow-speed shaker. For secondary antibody staining, HIOs were exposed to 80 μL of 200x-diluted, fluorophore-conjugated secondary antibody in the blocking buffer for 1 day at 4°C or 2 - 4 hours at room temperature on the slow-speed shaker. To remove unbound secondary antibody and stain with DAPI, HIOs were stained with 80 μL of 1 μg/mL DAPI in 1x PBS with 0.5% Triton X-100 for 25 minutes followed by washing with 80 μL of 1x PBS for 15 - 20 minutes at room temperature on the slow-speed shaker. HIOs were cleared by incubating in 30 μL of 1x pre-warmed (at 37°C) RapiClear 1.52 (Sunjin Lab, Taiwan) or Visikol^®^ Histo-M™ (SKU: HM-30, Visikol) in a clear flat-bottom, 384-well plate at room temperature for 0.5 - 1 hour. The stained and cleared HIOs were inspected under the Nikon A1 Rsi confocal microscope.

### Dynamic secretion of ghrelin from HIOs in the 36PerfusionPlate

HIOs were generated using iPSC lines containing a neurogenin-3 inducible construct to increase differentiation of enteroendocrine cells as previously described^31^. HIOs were patterned distally to either jejunum-like or ilium-like phenotypes with minor modifications to the differentiation protocol as described in literature^50^. After 24-hour doxycycline treatment, HIOs were cultured in 24-well plates for an additional 5 days. On day 6, HIOs were mixed with undiluted Matrigel and manually loaded on the 36PillarPlate (6 μL per pillar) using a multichannel pipette. The 36PillarPlate was then inverted and combined with the 36PerfusionPlate and incubated at 37°C for 5 minutes for gelation before adding the Gut maturation medium with EGF. Following overnight culture in the 36PillarPlate/36PerfusionPlate, HIOs were rinsed several times with warm PBS to remove residual medium. HIOs were then subjected to nutrient challenges by first loading the UR with 600 μL of 3 mM glucose in Krebs-Ringer Bicarbonate (KRB). After 30 minutes, 100 μL was removed from the LR and then 100 μL of fresh 3 mM glucose in KRB was added to the UR to maintain a constant flow rate. This was repeated every 30 minutes for a total of 2 hours in 3 mM glucose. All KRB was aspirated after 2 hours, and the process was repeated with 20 mM glucose in KRB for 3 hours. All samples were immediately frozen at - 80°C until analysis. KRB consisting of 140 mM NaCl, 0.15 mM Na_2_HPO_4_, 5 mM NaHCO_3_, 1 mM MgSO_4_, 4.6 mM KCl, 2 mM CaCl_2_, 0.05% BSA, and 30 mM HEPES at pH 7.4 was prepared freshly the day of or before nutrient challenges. Undiluted sample buffer was assayed for ghrelin content using ghrelin human ELISA kit (ThermoFisher) according to manufacturer specifications.

### Calculation of the IC_50_ value

Since the background luminescence of completely dead cells (following treatment with 70% methanol for 1 hour) was negligible due to background subtraction, the percentage of live HLOs was calculated using the following equation:

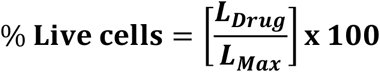

where L_Drug_ is the luminescence intensity of HLOs exposed to nefazodone and L_Max_ is the luminescence intensity of fully viable HLOs (control).

To produce a conventional sigmoidal dose-response curve with response values normalized to span the range from 0% to 100% plotted against the logarithm of test concentration, we normalized the luminescence intensities of all HLO spots with the luminescence intensity of a 100% live HLO spot (HLOs contacted with no compound) and converted the test compound concentration to their respective logarithms using Prism 8 (GraphPad Software, San Diego, CA, USA). The sigmoidal dose-response curve (variable slope) and IC_50_ value (i.e., concentration of nefazodone where 50% of HLO viability inhibited) were obtained using the following equation:

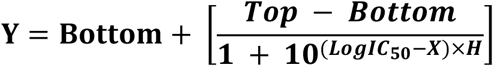

where IC_50_ is the midpoint of the curve, H is the hill slope, X is the logarithm of test concentration, and Y is the response (% live cells), starting from the top plateau (Top) of the sigmoidal curve to the bottom plateau (Bottom).

### Calculation of the coefficient of variation (CV)

To establish robustness of cell printing and spheroid transfer on the pillar plate, the range of error was measured using the coefficient of variation (CV). The CV value is the ratio of the standard deviation (SD) to the average (Avg). It represents variability in relation to the average signal strength, thus the inverse of the signal-to-noise ratio.

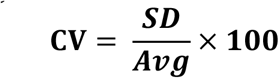

### List of antibodies used for cell staining

| Antigen | Host | Final concentration | Vendor | Catalog # |
| --- | --- | --- | --- | --- |
| Albumin | Rabbit | 10 µg/mL | Abcam | ab2406 |
| HNF4α | Mouse | 5 µg/mL | Santa Cruz | sc-374229 |
| E-cadherin | Mouse | 5 µg/mL | Santa Cruz | sc-21791 |
| CDX2 | Mouse | 10 µg/mL | ThermoFisher | 39-7800 |
| Alexa Fluor 488 donkey anti-mouse (H+L) IgG | Donkey | 1 - 10 µg/mL | ThermoFisher | A-21202 |
| Alexa Fluor 647 donkey anti-rabbit (H+L) IgG | Donkey | 1 - 10 µg/mL | ThermoFisher | A-31573 |

## Data availability

The authors declare that all the data supporting the results in this study are available within the article and its Supplementary Information.

## Acknowledgements

This study was supported by the National Institutes of Health (NIDDK UH3DK119982, NCATS R44TR003491, and NIEHS R01ES025779) and the Ohio Third Frontier Commission (TVSF Phase IB and Phase II).

## Author contributions

S.Y.K.: Tested the injection-molded pillar plate, developed surface chemistry of the pillar plate, performed flow pattern simulation with SolidWorks, tested flow rates within the perfusion well plate, performed the intestine organoid experiments, interpreted the results, and wrote the result and method sections.

M.K.: Performed the liver organoid experiments, interpreted the results, and wrote the result and method sections.

S.S.: Developed surface chemistry of the pillar plate, performed the liver and intestine organoid printing experiments, interpreted the results, and wrote the result and method sections.

P.L.: Performed dynamic hormone secretion experiments with intestine organoids, interpreted the results, and wrote the result and method sections.

S.J.L.: Tested the injection-molded perfusion well plate, developed surface chemistry of the perfusion well plate, interpreted the results, and wrote the result and method sections.

Y.C.: Performed the liver organoid experiments.

P.J.: Developed surface chemistry of the pillar plate and established spheroid printing protocols.

P.A.: Developed surface chemistry of the pillar plate.

J.L. and Y.Y.: Performed flow pattern simulation with SolidWorks.

J.G.S.: Cultured intestine organoids.

S.A.: Helped to write the introduction section.

E.A.: Helped to write the manuscript and supervised the project.

J.M.W.: Planned the intestine organoid experiments, interpreted the results, revised the manuscript, and supervised the project.

T.T.: Planned the liver organoid experiments, interpreted the results, revised the manuscript, and supervised the project.

M.Y.L.: Conceived and designed the pillar/perfusion well plates, developed surface chemistry, planned the bioprinting experiments for organoid development, interpreted the results, wrote and revised the manuscript, and supervised the project.

## Supplementary information

**Supplementary Figure 1.**
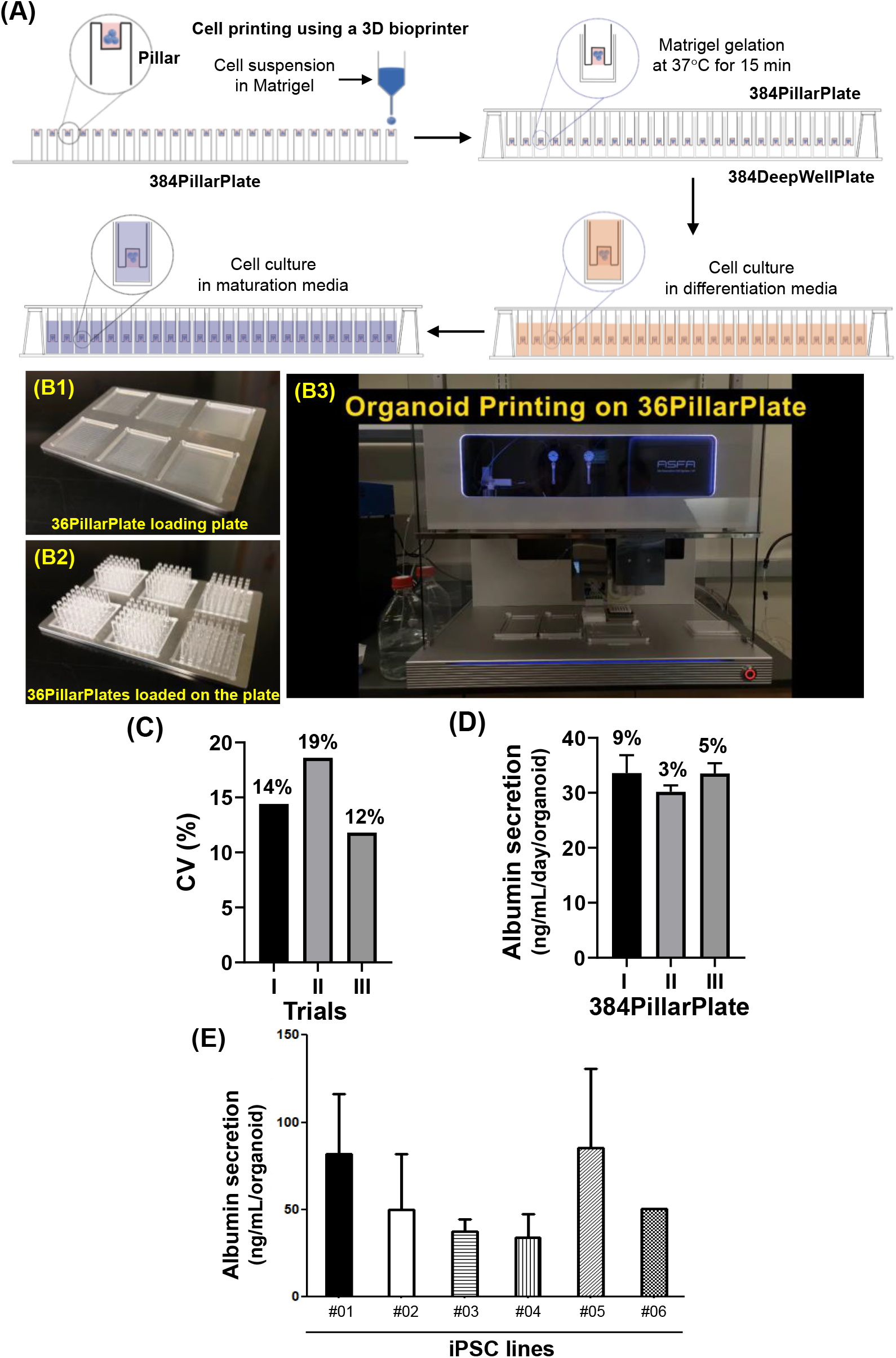
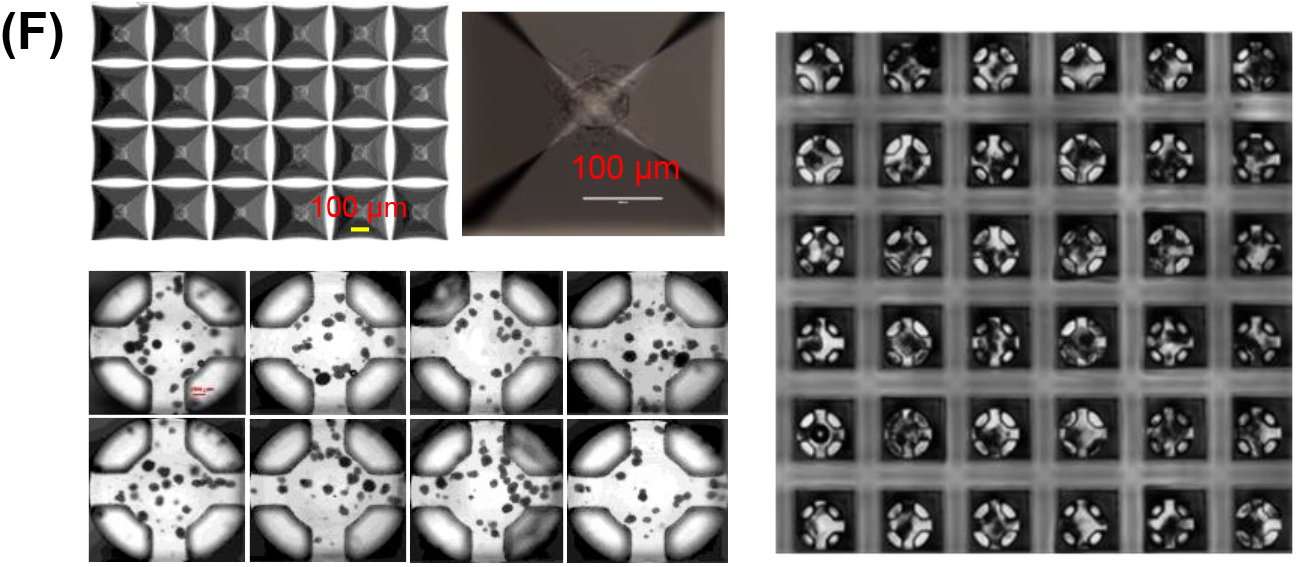
Cell printing strategy on the 36PillarPlate/384PillarPlate. **(A)** Schematic of printing single cell suspension in Matrigel on the pillar plates and organoid differentiation and maturation. **(B)** Cell printing on the 36PillarPlate: (B1) An aluminum plate made for cell printing on the 36PillarPlate, (B2) Six 36PillarPlates loaded on the aluminum plate, and (B3) Automatic cell printing on the 36PillarPlate using a 3D bioprinter (https://www.youtube.com/watch?v=ZiKvrAuFZbg). **(C)** The coefficient of variation (CV) for printing frozen foregut cells suspended in 2-fold diluted Matrigel on the 384PillarPlate. To evaluate the printing accuracy on the pillar plate, the CV values were calculated by measuring the viability of printed foregut cells in Matrigel using CellTiter-Glo^®^ luminescent cell viability assay kit. The low CV values for different trials indicate uniform cell printing on the pillar plate. **(D)** Albumin secretion (ng/mL/day/organoid) from HLOs on the 384PillarPlate to assess accuracy of printing frozen foregut cells in Matrigel. Albumin secretion from HLOs was measured on day 18 on the pillar plate (n = 9). The range of CV values obtained was 3.4 - 8.9%. **(E)** Measurement of albumin secretion from HLOs derived from multiple iPSC lines for day 25 on the 36PillarPlate by ELISA. Cell growth media containing albumin were collected from a 384-deep well plate after 24 h incubation, and albumin secretion was normalized to the number of HLOs (n = 3). **(F)** Accuracy of printing cell spheroids in Matrigel for organoid culture on the pillar plate: (**Upper panel**) Formation of iPSC spheroids in an AggreWell 400 plate. (**Lower panel**) Representative images of printed iPSC spheroids on the 36PillarPlate. Grey dots represent iPSC spheroids. iPSC spheroids from 3 wells (200 cells/microwell) were collected after 4 days of culture, centrifuged at 1,000 rpm for 1 min, resuspended in growth medium, mixed with GFR Matrigel at 1:1 ratio (3,000 spheroids per 400 μL), and printed on the 36PillarPlate (4 μL/pillar). Uniform spheroid printing was achieved repeatedly at high spheroid density. The CV value obtained from bioprinting experiments was 18.4%. The size of spheroids below 100 - 150 μm and high spheroid density were critical to maintain good suspension of spheroids in Matrigel and achieve high bioprinting accuracy. (**Right panel**) Stitched image of HIOs on the 36PillarPlate after 14 days of culture.

**Supplementary Figure 2.**
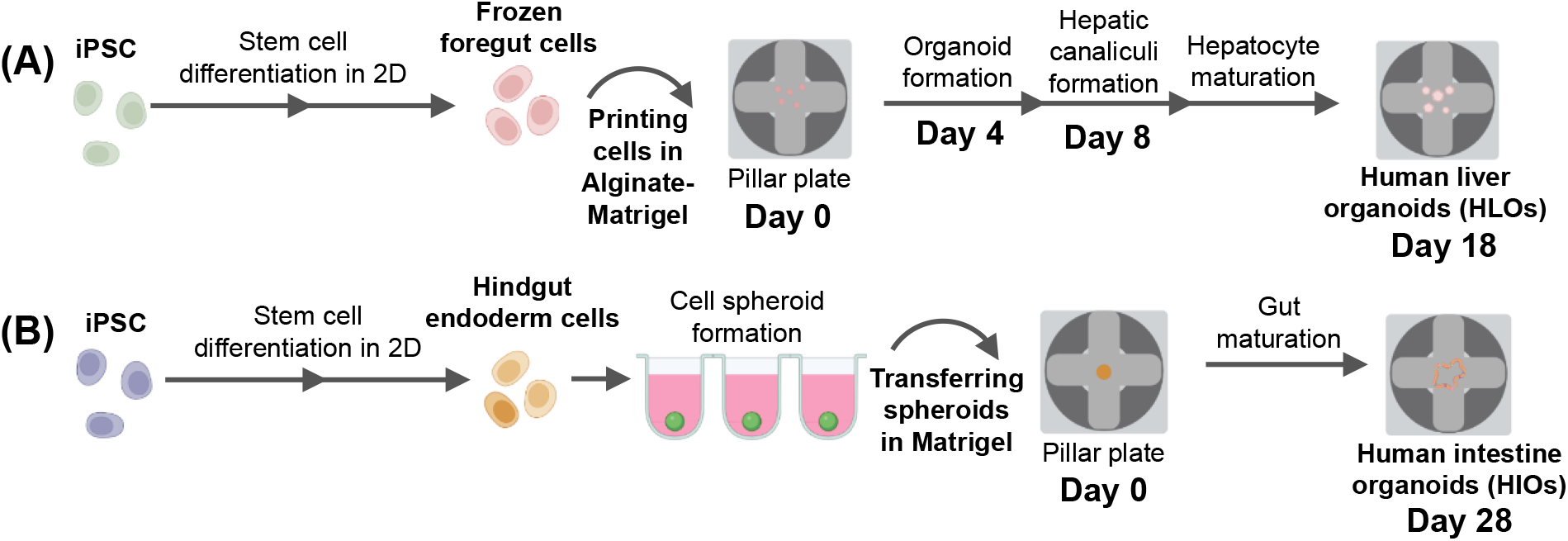
Steps necessary for organoid generation and timing considered for cell printing: **(A)** human liver organoid (HLO) generated by printing single cell suspension and **(B)** human intestine organoid (HIO) generated by transferring cell spheroids.

**Supplementary Figure 3.**
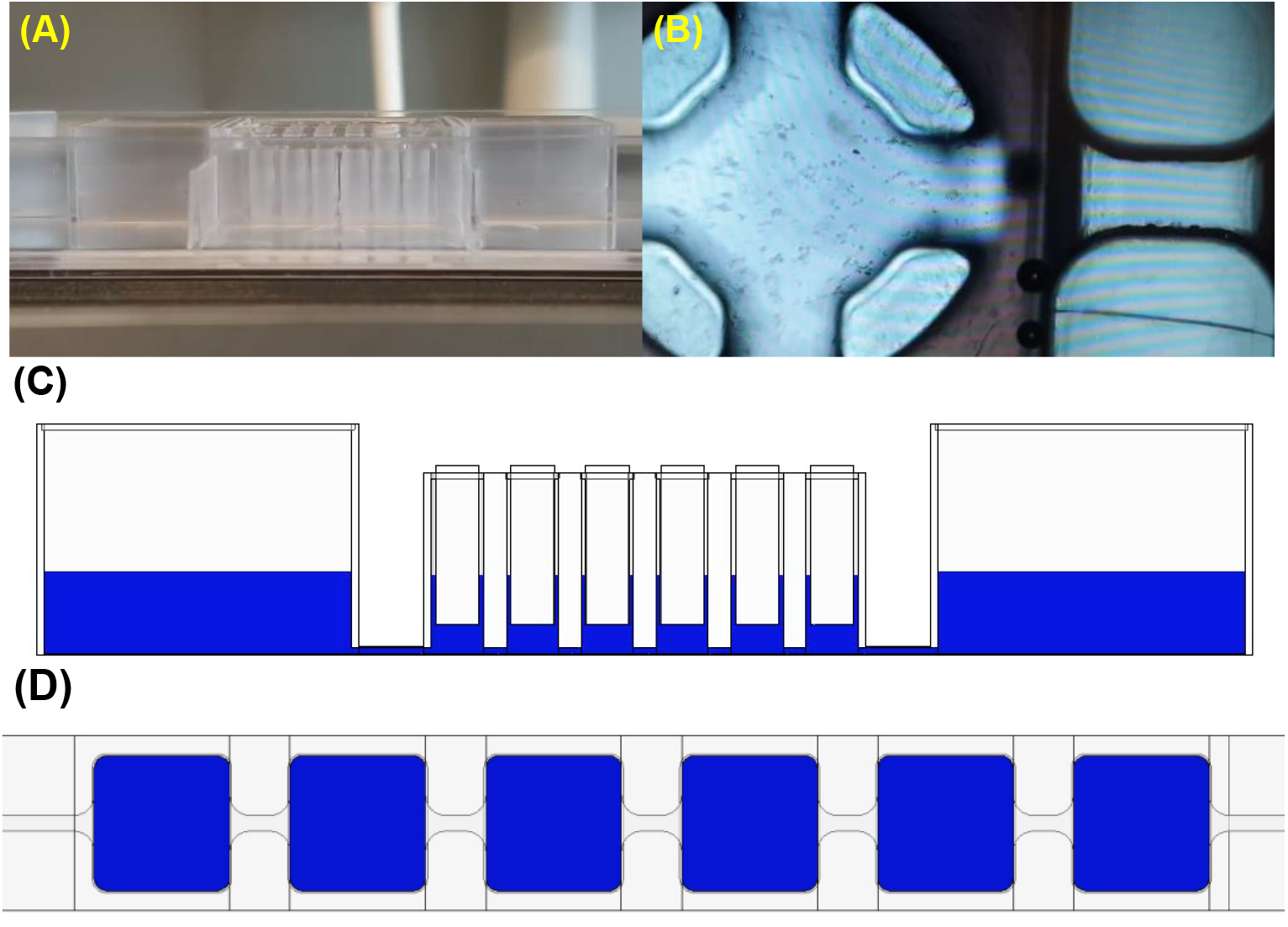
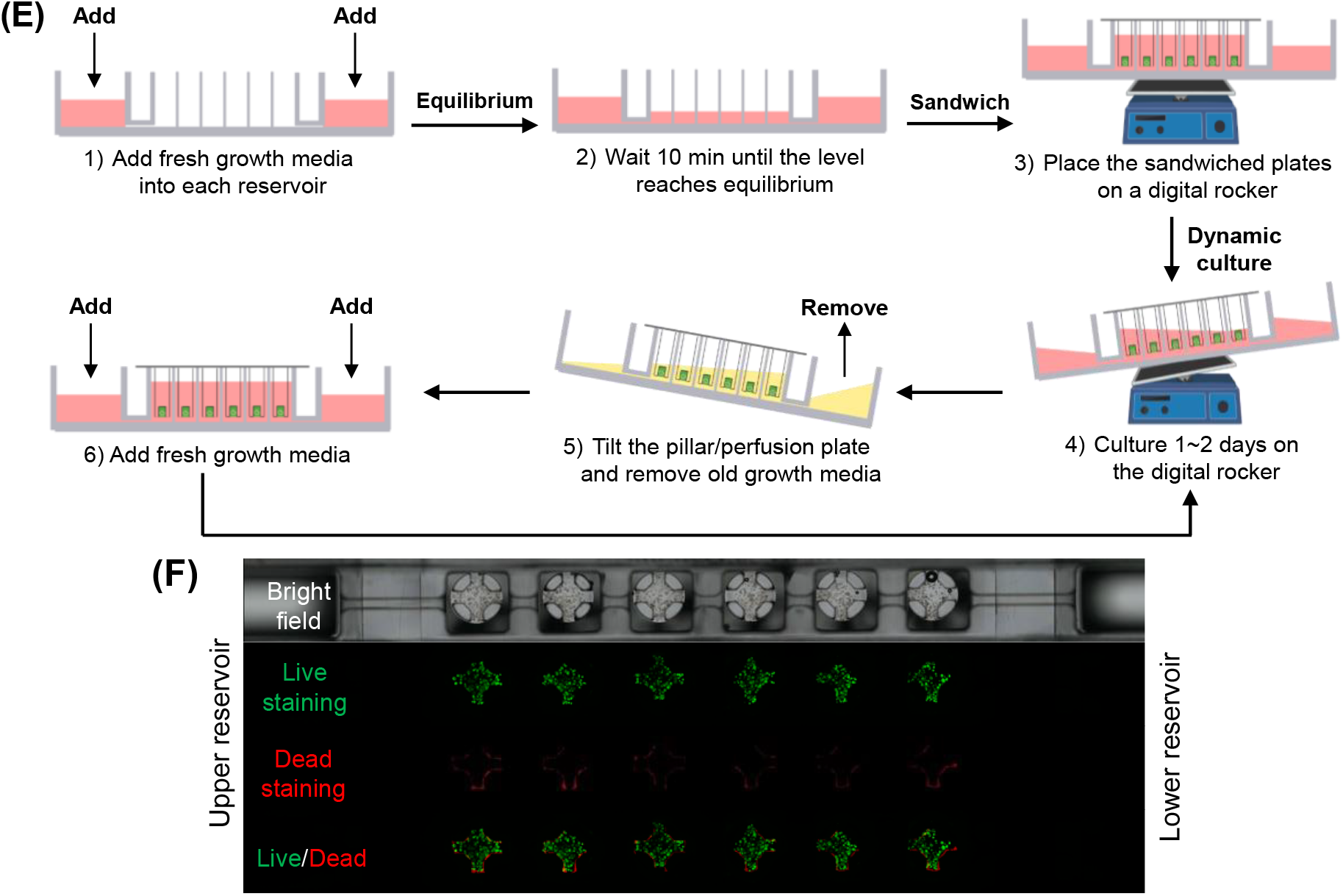
Experimental procedures with the 36PillarPlate and the 36PerfusionPlate. **(A)** Gravity-driven flow of trypan blue dye within the 36PerfusionPlate sandwiched with the 36PillarPlate (https://www.youtube.com/watch?v=Opsx-gdJKss). **(B)** Flow of trypan blue-stained Hep3B cells under the pillar through the microchannel in the 36PerfusionPlate (https://www.youtube.com/watch?v=odIsh27EJRg). **(C)** Changes in water levels in perfusion wells of the 36PerfusionPlate over time were simulated with SolidWorks with 900 μL of water at 10° tilting angle and 1 min frequency (https://www.youtube.com/watch?v=d_xVuK515S4). **(D)** Velocity profile under the pillars in the 36PerfusionPlate over time simulated with SolidWorks with 900 μL of water at 10° tilting angle and 1 min frequency (https://www.youtube.com/watch?v=gHIk9hWiNKw). **(E)** Experimental procedures for long-term dynamic organoid culture in the 36PerfusionPlate (https://www.youtube.com/watch?v=h-cs0Z2jzXE; https://www.youtube.com/watch?v=S0AWwhtoN2w). **(F)** Live/dead imaging of HLOs in the 36PerfusionPlate. HLOs on the 36PillarPlate were stained by perfusion flow of calcein AM and ethidium homodimer for 30 minutes in the 36PerfusionPlate.

